# Repeated evolution of supergenes on an ancient social chromosome

**DOI:** 10.1101/2024.12.01.626239

**Authors:** Aparna Lajmi, Pnina Cohen, Chih-Chi Lee, Zeev Frenkel, Yoann Pellen, Eyal Privman

## Abstract

Supergenes are non-recombining chromosomal regions that code for complex polymorphic traits. Advances in population genomics have uncovered supergenes associated with diverse traits, ranging from butterfly wing patterns to floral morphs. In ants, two supergenes on non-homologous “social chromosomes” in *Solenopsis* and *Formica* are associated with social polymorphism, with either single queen (monogyne) or multiple queens (polygyne) colonies. We discovered a new supergene associated with similar polymorphism in the desert ant *Cataglyphis niger*. Despite *Cataglyphis* being more closely related to *Formica* than *Solenopsis*, its social chromosome is homologous to that of *Solenopsis*, with conservation of synteny in gene content and order. This suggests that the social chromosome is ancient, dating back to the common ancestor of *Solenopsis* and *Cataglyphis*, at least 90 million years ago. Low sequence divergence between supergene haplotypes in both *Solenopsis* and *Cataglyphis* suggests that the two supergenes evolved recently and independently in the two divergent lineages on this ancient social chromosome. Comparative analysis of hymenopteran genomes further revealed a bee chromosome homologous to the ants’ social chromosome. The ant social chromosome contains the largest set of genes that are conserved as a linkage group across ant and bee genomes. This conserved gene set is enriched for olfactory functions, most notably a large number of odorant-binding proteins. The conservation of this gene set suggests that this chromosome plays an important role in social behavior across social Hymenoptera. We propose that the conserved gene set in the social chromosome was repeatedly used as a pre-adapted toolkit for the evolution of social traits in general, and specifically in the evolution of polygyne social structure in ants.

## 1. Introduction

Convergent evolution is a common phenomenon across the tree of life^1^, yet the genomic architectures that facilitate the repeated evolution of complex traits remain poorly understood. Social behavior and the underlying sociobiological traits, particularly in Hymenoptera (ants, bees, and wasps), exemplify complex phenotypes that have evolved repeatedly in multiple lineages. Previous literature suggests that the evolution of sociality in diverse hymenopteran lineages involved the reuse of a core set of conserved genes that were originally involved in other biological processes, a hypothesis termed the genetic “toolkit” hypothesis^2^. This hypothesis suggests that genes originally involved in solitary behaviors or other biological processes were co-opted to facilitate sociality, in line with the notion that evolutionary innovations often emerge from preexisting genetic frameworks. For example, the *foraging* gene that is conserved across insects and regulates foraging behavior in *Drosophila melanogaster*^3^ has been co-opted in bees and ants to regulate caste-specific behaviors^4,5^. Foraging and brood care behaviors in the worker caste are key to the success of social insect colonies. Maternal care, which is not common in insects but found in many solitary wasps, has been proposed to be a pre-adaptation for the evolution of sociality^6,7^. Such pre-adaptations in solitary Hymenoptera may have facilitated the repeated evolution of sociobiological traits in this clade.

Ants have repeatedly evolved the trait of multiple-queen colonies (polygyny) from an ancestral phenotype of a single-queen colony (monogyny)^8,9^. Polygyny is a complex trait involving a suite of life-history, behavioral, physiological, and morphological modifications^10,11,12^. Moreover, it results in nestmates that have very low relatedness, which undermines the evolution of cooperation by means of kin selection^9,12^. Hughes *et al.*^8^ reported that polygyne colonies were observed for 54 out of 130 (42%) ant species included in that study. Some species of ants are socially polymorphic, where both monogyne and polygyne colonies coexist in the same population. A genetic basis of this social polymorphism has been documented and examined in detail in two distantly related ant lineages, *Solenopsis* fire ants and *Formica* ants. In both lineages, a supergene was found to be associated with this polymorphism^13^. These supergenes are chromosomal regions with suppressed recombination, containing multiple tightly linked loci that code for a complex trait. The most well-known type of supergenes is sex chromosomes, but virtually any other complex polymorphic trait may be coded by a supergene^14^. Further studies have identified additional species in the *Solenopsis* and *Formica* lineages that have homologous supergenes. Recent evidence for additional non-homologous supergenes for social polymorphism has been reported in two other lineages, *Pogonomyrmex* and *Myrmica*^15,16^. Social supergenes in some of these species are associated with additional phenotypic diversity beyond colony queen number, including queen miniaturization^17^ and split sex ratio^18^. The rapid increase in the discovery of ant lineages with social supergenes offer an opportunity to investigate the genomic basis of the recurrent evolution of complex traits.

The two well described social supergenes in *Solenopsis* and *Formica* showcase several important similarities and differences in how the supergene is maintained. The first social supergene was discovered by Wang, Wurm, and colleagues in the red fire ant *Solenopsis invicta*, which is native to tropical South America^19^. In monogyne colonies of this species, the queen and the workers are MM homozygotes at the supergene (the *Solenopsis* supergene haplotypes were originally named *SB* and *Sb*, but we are using M for monogyne-associated and P for polygyne-associated haplotypes across all species for consistency). On the other hand, the polygyne queens are always MP heterozygous at the supergene (with very rare exceptions), and produce workers of all three genotypes: MM, MP, and rarely PP, by mating with both M and P haploid males^20^. The almost complete absence of the PP genotype in queens led the authors to refer to this chromosome as a “Y-like social chromosome”. This supergene was found across a few closely related, socially polymorphic *Solenopsis* species^21–23^ that speciated approximately one million years ago (MYA)^24^. The formation of the supergene was dated approximately to the same time based on the divergence of the M and P haplotype sequences^24,25^. In the Alpine silver ant *Formica selysi*, where the second social supergene was discovered, monogyne queens and workers are MM homozygotes at the supergene while the polygyne queens and workers are either PP homozygotes or MP heterozygotes^26^. Unlike in *S. invicta*, MM genotypes in polygyne colonies are eliminated in this species, which was suggested to be due to a maternal effect killing^27^. Supergenes homologous to the *F. selysi* supergene were found in numerous *Formica* species and therefore evolved in their common ancestor at least 30 MYA^28,29^. Recent studies reveal considerable diversity of the supergene system in diverse *Formica* species, including genomic and phenotypic variations^16–18^. *Formica* and *Solenopsis* belong to two different ant subfamilies (Formicinae and Myrmicinae) that diverged more than 90 MYA^30^. The young age of the two supergenes relative to the lineage divergence and the fact that they reside on non-homologous chromosomes imply that the two supergenes have independent origins.

The desert ant *Cataglyphis niger* is a socially polymorphic species, with populations consisting of both monogyne and polygyne colonies. These populations were initially thought to consist of multiple species, based on divergent mitochondrial sequences (mitotypes) that are associated with the social form^31^. However, nuclear genomic sequencing revealed no genetic divergence between monogyne and polygyne colonies, demonstrating that *C. niger* is a socially polymorphic species^32^. The genera *Cataglyphis* and *Formica* belong to the same tribe Formicini within the Formicinae subfamily, which diverged approximately 40 MYA^30^. Therefore, *C. niger* is an interesting target species for the study of the evolution of social polymorphism across the ant phylogeny.

Here, we examined the genomic basis of social polymorphism in *C. niger* by generating reduced representation genome sequencing data for 672 individuals from 30 nests representing both social forms from a single population. The social form of the nests was ascertained by estimating relatedness among nestmates, as well as other lines of evidence, including aggressive behavior between colonies. We discovered a novel “Y-like” social chromosome in *C. niger* harboring a supergene associated with the social polymorphism. We find complete linkage of the P haplotype with the maternal lineage, indicating that it is strictly maternally inherited. Surprisingly, we find that the *Cataglyphis* social chromosome is homologous to the *Solenopsis* social chromosome, even though the two lineages diverged more than 90 MYA. A wider survey across multiple hymenopteran genomes revealed conserved synteny of the social chromosome across bees and ants. Our study revealed a conserved gene set on an ancient chromosome that was repeatedly used in the evolution of supergenes coding for alternative social structures in diverged ant lineages. We hypothesize that this gene set represents a genetic “toolkit” that may have been involved in the evolution of various sociobiological traits in Hymenoptera. This gene set evolved in a common ancestor of ants and bees, where it may have originally coded for a different trait. This ancestral trait may have been a pre-adaptation for sociality, such as regulation of maternal care.

## 2. Results and discussion

### 2.1 Social polymorphism in *Cataglyphis niger* is associated with chromosome 2

We fully excavated 30 nests from a single population in Tel Baruch, Israel, where both monogyne and polygyne nests were known to co-exist. Genomic data was generated for 672 samples (28 queens and 644 workers) from the 30 nests (20-24 individuals each) using double-digest restriction-site associated DNA sequencing (RAD-seq), yielding genotype data for 30,955 high-confidence single nucleotide polymorphisms (SNPs). As queens were found only in 14 nests (7 had one queen and 7 had more than one queen), the social form of the sampled nests was assigned based on the relatedness within the nest.

A pairwise kinship matrix constructed based on the total length of shared identity-by-descent (IBD) haplotypes across 672 sampled individuals showed some nests with relatedness between all nestmates and some where most pairs were unrelated (Figure 1a), as expected for monogyne and polygyne nests, respectively. In monogyne nests, we expected three clusters of relatedness scores— full siblings or super sisters sharing 75% of their genome, mother-daughter pairs with 50% relatedness, and half-sisters (from different fathers) with 25% relatedness. The nests with higher relatedness had broadly three clusters of relatedness scores consistent with our expectations in a monogyne nest (Figure 1b and Supplementary Figure S1a). However, we did find a small number of worker pairs with 50% relatedness, which could result from the queen mating with brothers. As expected for polygyne nests, most pairs of workers had relatedness scores lower than half-sister relatedness (25%). Based on the distribution of kinship scores, 13 nests were classified as monogyne and 17 as polygyne for downstream analyses (Supplementary Figure S1b and c). A one-way ANOVA showed that aggression scores differed significantly among colony types, *F*(2, 101) = 40.73, *p* < .001, η² = 0.45. Post-hoc tukey tests showed higher aggression between monogyne– polygyne/monogyne–monogyne nests than controls (mean diff = 0.73, *p* < 0.001) and lower aggression between polygyne nests than controls (mean diff = −0.53, *p* < 0.001), with no difference between polygyne nests and controls (*p* = 0.116). The number of queens found during sampling was also in line with these results (full details provided in Supplementary Figures S2 & S3).

**Figure 1:**
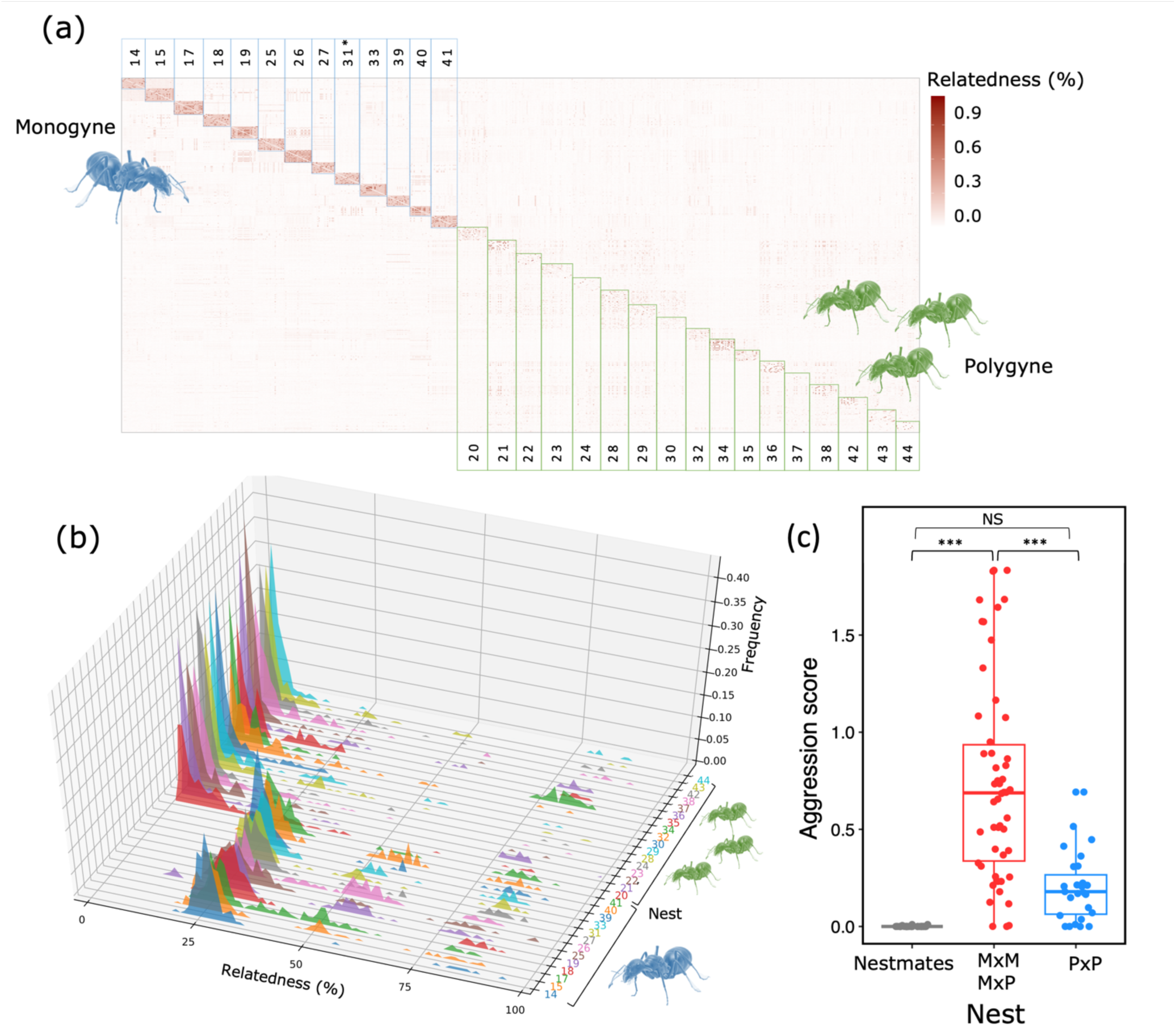
Identification of social form using kinship inference based on the total length of identity-by-descent (IBD) haplotypes shared between nestmates, **(a)** plotted as a kinship matrix demarketing monogyne nests in blue and polygyne nests in green. **(b)** Distributions of relatedness within nests. **(c)** Aggression scores from the dyadic assay showed significantly higher aggression between monogyne and monogyne-polygyne nests compared to between polygyne nests. Stars indicate *p-*value of < 0.001

We examined *F_ST_* at each SNP between monogyne and polygyne samples (Figure 2a). The average weighted *F_ST_* across the genome was low (0.019) as expected due to gene flow and lack of genetic differentiation between the two social forms^32^. However, the average weighted *F_ST_* across chromosome 2 was much higher (0.12) than the rest of the genome. 164 high *F_ST_* SNPs (>0.2) were found across a ∼5.8Mb stretch of chromosome 2, from position 3,930,062 bp to 9,747,310 bp in the Cnig_gn3.1 genome assembly. These 164 SNPs represent only 0.15% out of the 112Kb that were covered by our RAD-seq data in the supergene region (with depth ≥ 12), giving an estimate of the average sequence divergence between the M and P haplotypes. This rather low sequence divergence is surprisingly similar to the 0.14% divergence reported in *Solenopsis* (15,367 fixed SNPs across the 10.8 Mb supergene region^33^). The low sequence divergence was used to date the *Solenopsis* supergene to 1.1 (0.7-1.6) MYA^25^. While we do not have comparable whole-genome sequence data in *Cataglyphis* to date the supergene with the same phylogenetic approach and the same level of accuracy as was done in *Solenopsis*, we can conclude that the age of the *Cataglyphis* supergene is on the same order of magnitude.

**Figure 2:**
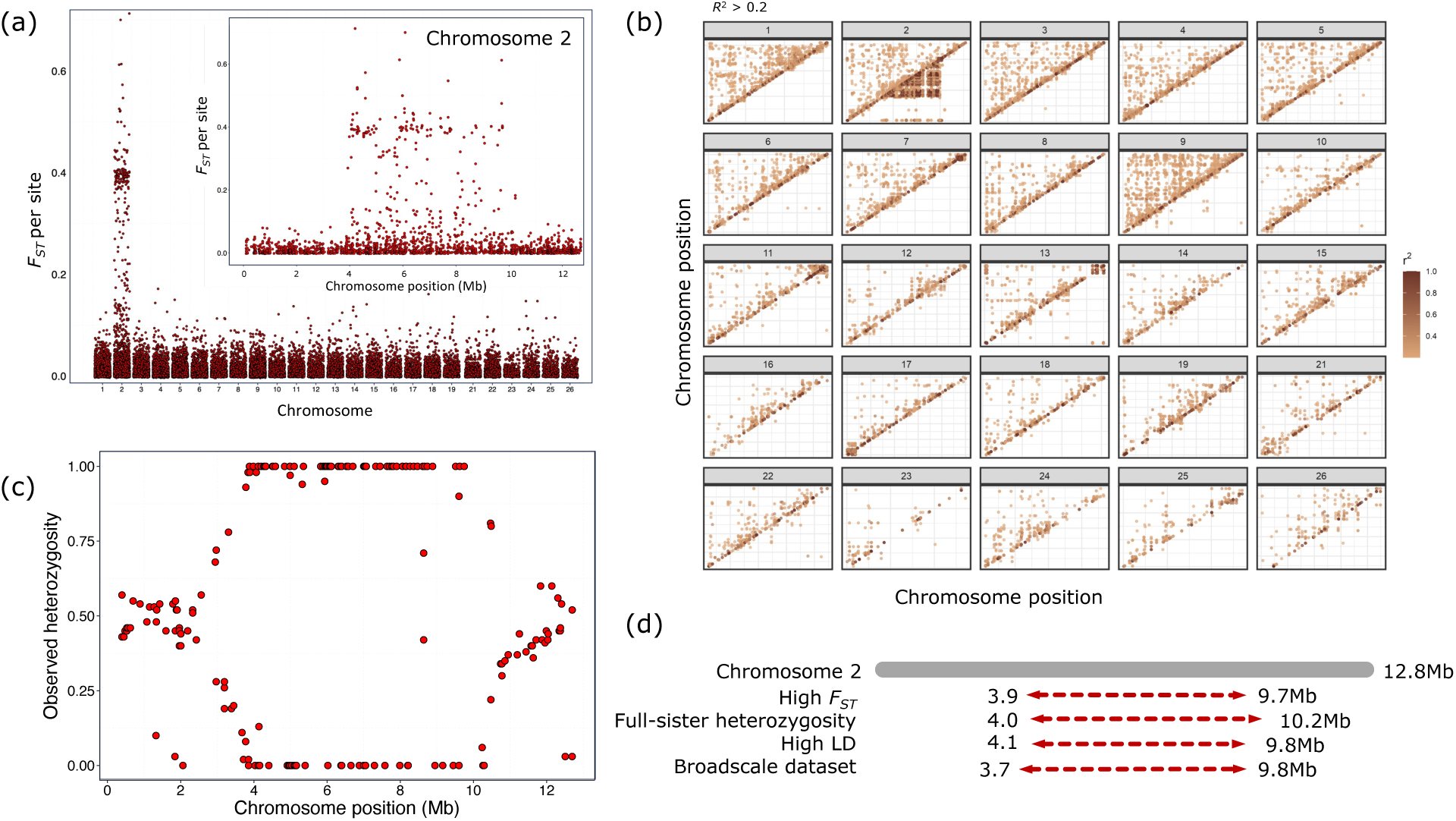
Genetic divergence and high linkage disequilibrium on chromosome 2 suggest the presence of a supergene. **(a)** Plot of *FST* between social forms (defined based on kinship) showing high *FST* values on chromosome 2 . **(b)** Heatmaps of linkage disequilibrium in monogyne (above the diagonal) and polygyne (below the diagonal) population samples (displaying only *r*^2^>0.2) across all chromosomes. **(c)** Heterozygosity along chromosome 2 in 69 full sisters from a polygyne colony. **(d)** Comparison of the boundaries of the supergene region using different criteria.

### 2.2 Suppression of recombination in chromosome 2

Supergenes maintain complex polymorphic traits despite high gene flow between morphs by suppression of recombination that prevents the shuffling of the alleles associated with the trait at different loci. We tested for suppression of recombination using two approaches: (1) testing for linkage disequilibrium (LD) in the population sample and (2) testing for linkage in progeny of a single polygyne queen. We conducted LD analyses separately on monogyne and polygyne samples to examine if the supergene region on chromosome 2 showed suppression of recombination. The analysis revealed high LD (*r*^2^ ≥ 0.2) in polygynes but not in monogynes across most SNP pairs in the supergene region (chromosome 2 positions 3,813,975 – 10,147,746 bp), aligning with the region of high *F_ST_* SNPs (Figure 2b). We also noted LD decaying rapidly with SNP distance across all chromosomes of monogyne and polygyne samples except in chromosome 2 in polygynes, where LD did not decay over long distances (Supplementary Figure S4). Adjacent to the large 5.7Mb LD block, a smaller 270 kb LD block can be observed in both monogyne and polygyne samples (Supplementary Figure S4). Similarly, four additional small LD blocks spanning over 249 kb were observed in other chromosomes of both the social forms (chr07: 298 kb; chr11: 249 kb; chr13: 434 kb; chr17: 297 kb) (Figure 2b). Since there is no linkage disequilibrium between these LD blocks and chromosome 2, and no association between loci in these genomic regions and the social form, these LD blocks are unlikely to be involved in maintaining the social polymorphism.

Additionally, linkage analysis was conducted based on RAD sequencing of 225 brood samples collected from a single polygyne queen. The mother was also sequenced. Six patrilines were identified in this data set and the largest patriline (69 full sisters) was used to build a linkage map. However, the genotype data did not allow for building a map of chromosome 2 because of an unusual pattern. All sisters had the same genotypes across the supergene region: 131 SNPs (70%) were always heterozygous, and 44 SNPs (24%) always had the same homozygous genotype, out of 186 SNPs that were heterozygous in the mother (Supplementary Figure S6). Plotting heterozygosity along chromosome 2 (Figure 2c) showed that SNPs in the supergene region were either 100% or 0% heterozygous. The extent of markers with heterozygosity of 100% spanned chromosome position 3,986,390 - 10,237,665bp. This pattern can be explained by a combination of two effects: (1) there were no recombination events within the supergene region, and (2) all the daughters received the same haplotype of the supergene from their mother. That is, recombination in the supergene was fully suppressed and one of the two maternal haplotypes is lethal. We note that this pattern is reminiscent of maternal effect killing of homozygous MM daughters in polygyne *F. selysi* colonies.

### 2.3 Maternally inherited ‘Y-like’ supergene

We examined the genotypes of 164 high *FST* (>0.2) SNPs on chromosome 2 by generating a heatmap (Figure 3a). The heatmap showed that polygyne samples were predominantly heterozygous (127/164 loci on average), whereas monogyne samples were largely homozygous for one allele (132/164 loci on average). This suggests that the polygyne social form is associated with a ‘Y-like’ supergene haplotype, which we denote as the P haplotype. That is, polygyne samples carry a heterozygous MP supergene genotype, while monogyne samples are MM homozygotes. This pattern holds for all queens and most of the workers. One conspicuous exception to this rule was nest M31, which was designated as a monogyne nest based on high relatedness, and yet, all samples in this nest had the MP genotype. The phenotype of high relatedness in this nest could have been observed if polygyne nests in *Cataglyphis* were also founded independently by queens and not, as commonly assumed, through nest budding. We speculate that some polygyne nests may be founded by a single MP queen, which is subsequently joined by additional MP queens. Indeed, this nest had a relatively small number of workers, as expected for a newly established nest. This is also in line with the geographic location of this nest among other polygyne nests (Figure 3c). The mitotype (discussed below) and aggression scores of this nest are also as expected for the polygyne form. The aggression scores of this nest against polygyne nests (Supplementary Figure S2) were relatively low (0.2 – 0.9) compared to its score against a monogyne nest (1.3). There were seven other exceptions to the broad pattern observed in the heat map, all of which were workers: five samples from polygyne nests had an MM genotype (four from P34 and one from P35) and two samples from monogyne nests had an MP genotype (M19-44 and M27-19). Nest details along with supergene genotypes and mitotypes are given in Supplementary Table S1.

**Figure 3:**
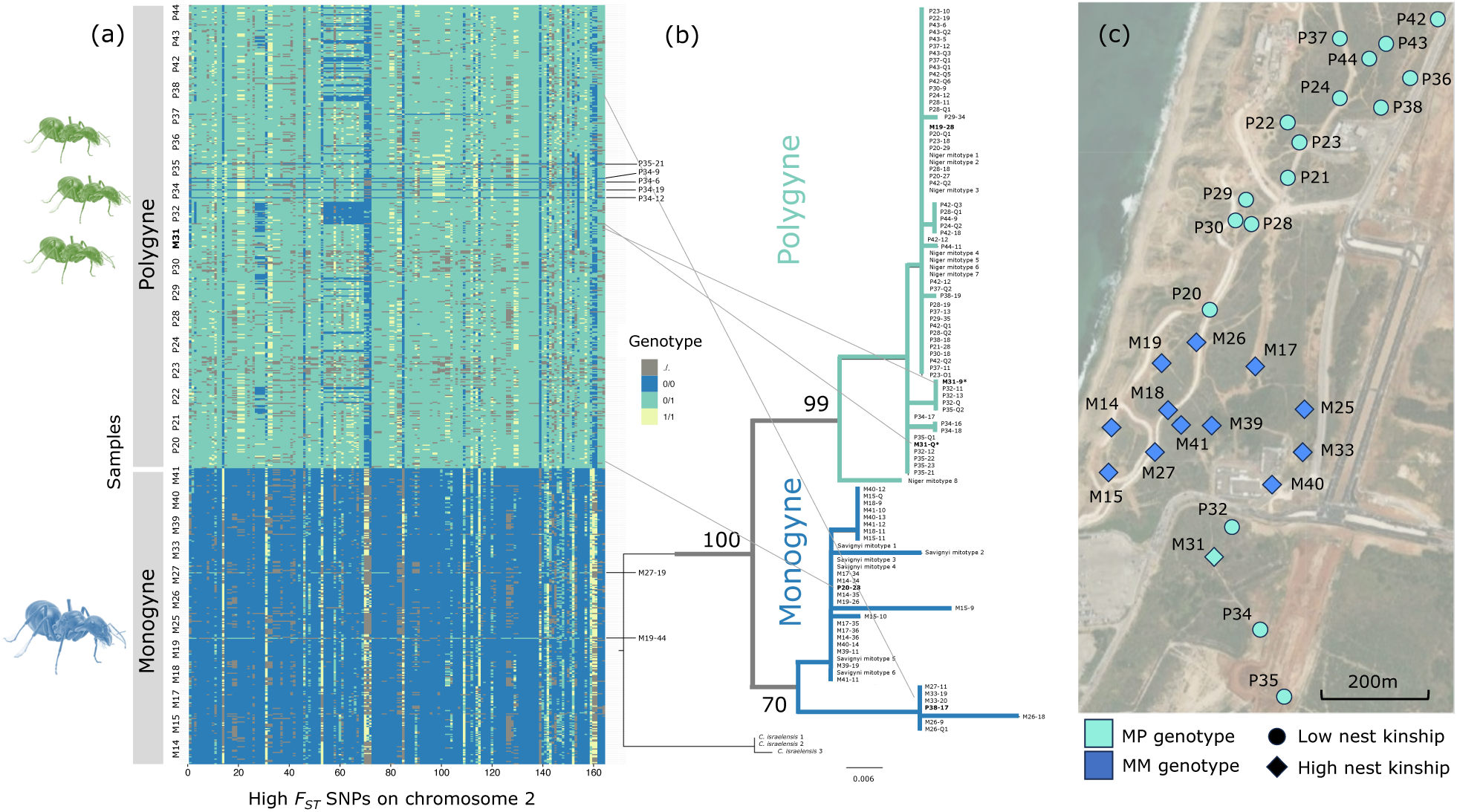
**(a)** Heatmap showing the genotypes at SNPs with high *FST* (>0.2) on chromosome 2. Generally, one allele is only found in heterozygous genotypes in polygyne colonies, whereas the other allele is homozygous in monogyne colonies. Exceptions to the rule are highlighted in nests M19, M27, M31, P34. **(b)** Mitochondrial phylogeny with a deep divergence between monogyne and polygyne samples indicating maternal inheritance of the social chromosome. **(c)** Geographic location of the nests where the social form was identified based on kinship and supergene genotype.

Previous work on this species suggested deep divergence between mitotypes associated with monogyne and polygyne social forms based on samples collected across Israel^31^. To ascertain if this pattern was consistent across the nests sampled in this study, we sequenced part of the mitochondrial gene Cytochrome *b* from two to three individuals from each nest. The mitochondrial phylogeny (Figure 3b) resulted in two deeply divergent well-supported monophyletic clades largely corresponding to monogyne and polygyne samples. Previously published mitotypes associated with monogyne nests (previously named the “savignyi” and “drusus” mitotypes) were nested within the monogyne clade and mitotypes associated with polygyne nests (the “niger” mitotype) were nested within the polygyne clade. This divergence in mitotypes associated with social form indicates a strict maternal inheritance of the social form.

Five samples were an exception to this rule. Both samples sequenced from nest M31 (a queen and a worker), which had the MP supergene genotype, also carried a polygyne-associated mitotype. Two worker samples from polygyne nests with MP supergene genotypes carried a monogyne-associated mitotype.

One conspicuous result from the heat map is the absence of both homozygous PP and MM genotypes from polygyne nests, where virtually all individuals have a heterozygous MP genotype. We note that while our sample consists of hundreds of workers from 18 polygyne nests, it only represents a single study site, and other polygyne populations in the species may differ. Nevertheless, within the study population, the large sample size makes the absence of homozygous genotypes highly conspicuous. Interestingly, this unexpected genotype distribution closely resembles polygyne colonies from the species *Formica francoeuri*^34^. The absence of MM genotypes suggests negative selection, possibly a result of mechanisms such as meiotic drive, maternal effect killer, or green beard effect, which were suggested for the social supergenes in other species.

Generally, it was suggested that the P chromosomes are selfish genetic elements. In *Formica selysi*^27^, the P chromosome is implicated in maternal effect killing, whereby the presence of the P chromosome in the mother results in the death of progeny that did not inherit at least one P chromosome, and in *Solenopsis invicta*^35^, it is implicated in causing a green beard effect—killing of queens that do not have the P chromosome.

Selfish mechanisms may be generally responsible for the stability of these social supergenes. The absence of PP genotypes also indicates negative selection, potentially due to deleterious mutations, which may be similar to the low fitness of PP queens in *S. invicta*. Such deleterious mutations may also result in P being lethal in males or in P males being sterile (although this is not the case in *S. invicta*). If so, MP daughters would always inherit their P chromosome from their mother (and M from their father), in line with the mitochondrial phylogeny. Another possible explanation for the maternal inheritance of the P chromosome is that P males do exist, but monogyne queens do not mate with them or have extremely low sperm transmission, as observed in *S. invicta*^36^.

### 2.4 The supergene is distributed across multiple populations in Israel

We used a previously published RAD-seq dataset of worker samples collected across Israel to verify that the supergene is distributed beyond our study population of Tel Baruch^32^. This study used different enzymes from our study to generate genomic libraries resulting in a different set of genomic loci. We used the mitotype of the samples as a proxy to identify their social form and calculated *FST* between inferred monogyne and polygyne samples at each SNP across the genome. High *FST* SNPs were detected on chromosome 2 between positions 3,985,614-9,804,010, which overlapped the supergene region identified in the Tel Baruch population (Supplementary Figures S7a and S6b). Similar to the pattern reported above (Figure 3a), the 164 high *FST* SNPs (>0.4) in this dataset were mostly homozygous MM (52/56) in the monogyne mitotype samples and heterozygous MP (13/14) in the polygyne mitotype samples (Supplementary Figure S7c). This sampling across Israel shows that most of the range consists of pure MM populations, whereas the P haplotypes are always found in polymorphic populations (MM and MP) limited to the central coastline region between Ashkelon and Netanya (Supplementary Figure S7d).

### 2.5 Repeated evolution of social supergenes on an ancient chromosome

Non-homologous social chromosomes are known to be associated with social polymorphism in *Solenopsis* and *Formica*. We wanted to examine whether the *Cataglyphis* social chromosome is homologous to either of these chromosomes. *Cataglyphis* and *Formica* both belong to the subfamily Formicinae, whereas *Solenopsis* belongs to the subfamily Myrmicinae. The two subfamilies diverged around 93 MYA^30^ (95% credible interval 91–95 MYA). Synteny dot plots based on one-to-one orthologs showed that the social chromosomes of *Cataglyphis* and *Formica* were not homologous (Figure 4a). Surprisingly, despite belonging to different subfamilies, the social chromosomes of *Cataglyphis* and *Solenopsis* are homologous (Figure 4b). Synteny is not only conserved in terms of gene content (macrosynteny) but also gene order (microsynteny), especially in the supergene region (red points in Figure 4c). Interestingly, the largest region with conserved gene order is inside both supergenes, while the non-overlapping regions of the supergenes show conservation of gene order to a lesser extent.

**Figure 4:**
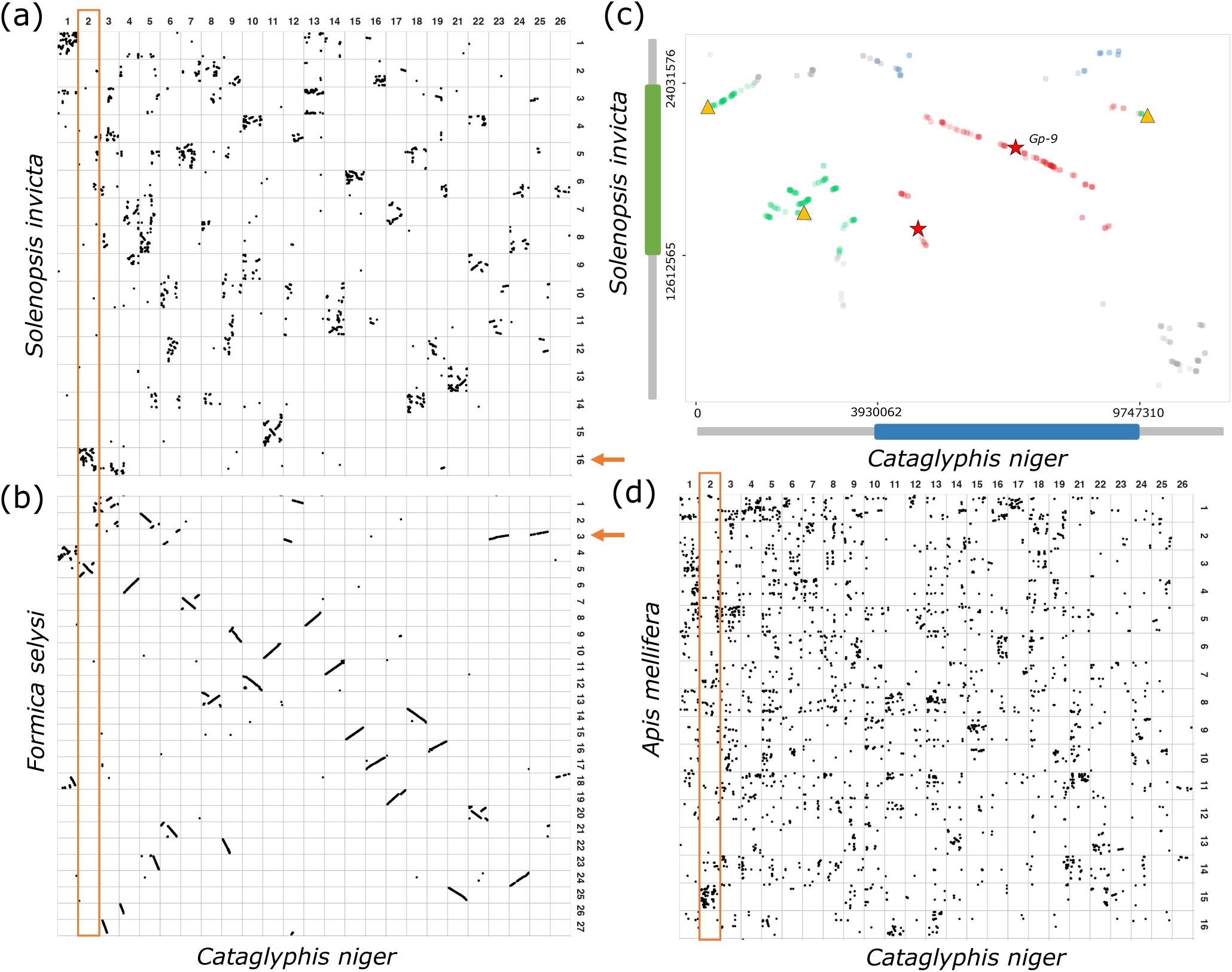
Conserved synteny of the social chromosome across ants and bees. Dot plots of one-to-one orthologs **(a)** between *Cataglyphis* and *Solenopsis* and **(b)** between *Cataglyphis* and *Formica*. Orange arrows indicate the social chromosome in *Solenopsis* and *Formica*, and the orange box indicates the social chromosome in *Cataglyphis*. **(c)** Social chromosomes with the supergene regions of *Cataglyphis* in blue and *Solenopsis* in green. Supergene coordinates in *Cataglyphis* are defined based on the high *FST* region (3,930,062 to 9,747,310bp), and in *Solenopsis* based on the inversion breakpoints (Yan *et al.*^37^). The overlapping supergene region highlighted in red shows remarkable conservation in gene order. The yellow triangles indicate odorant receptors (ORs) and red stars indicate odorant-binding proteins (OBPs). **(d)** *Cataglyphis* and *Apis* dot plot. *Cataglyphis* chromosome 2 shows notable conservation of gene content and homology to *Apis* chromosome 15.

To investigate the extent of the conservation of synteny of the social chromosome, we conducted a comparative genomic analysis of all published annotated, chromosome- level ant assemblies. We also added representatives of bees and wasps to this analysis amounting to 22 chromosome-level assemblies that allowed us to evaluate the conservation of synteny more broadly across Hymenoptera. As an estimate of macrosynteny, we calculated the percentage of *Cataglyphis* genes found on the homologous chromosome (the chromosome carrying the largest number of homologous genes) in each of these species (one-to-one orthologs only). Figure 5a shows that *Cataglyphis* chromosome 2 is one of the more highly conserved chromosomes across Hymenoptera, especially relative to other large chromosomes (chromosomes are numbered from largest to smallest). This chromosome is notably more conserved than most other chromosomes in distant lineages such as *Solenopsis* (75% of one-to-one orthologs on the same chromosome) and even in the honey bee *Apis mellifera* (41%).

**Figure 5:**
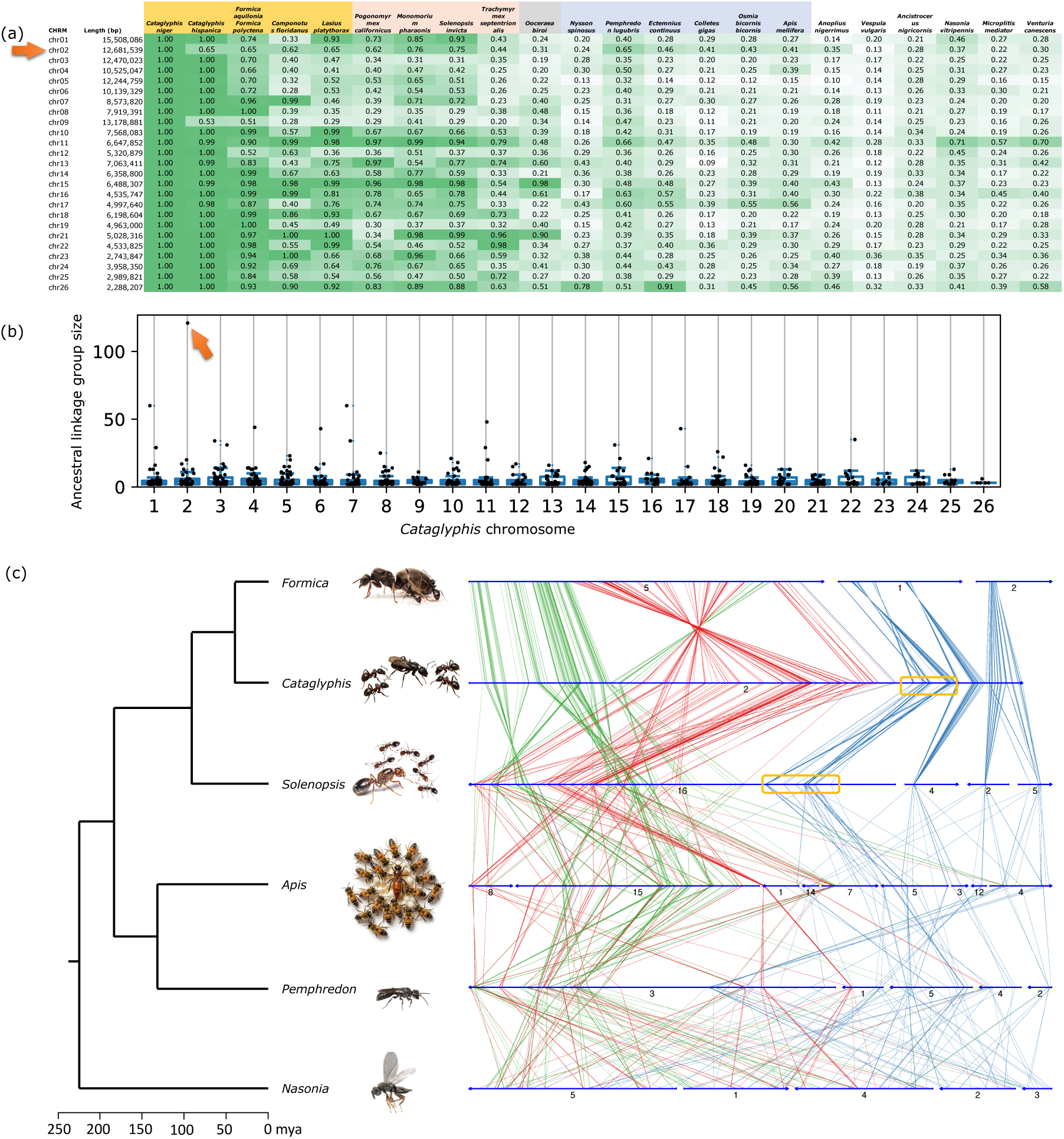
Conservation of synteny of all *Cataglyphis* chromosomes. **(a)** Percent of one-to-one orthologous genes in each *Cataglyphis* chromosome that reside in one chromosome in other Hymenoptera species. **(b)** The number of one-to-one orthologous genes in Ancestral Linkage Groups (ALG) of ants and bees. The star marks the largest ALG with 121 genes. **(c)** Synteny plot showing three representative species of ants, two Apoidea species, and one wasp outgroup. The figure highlights three broad regions of *Cataglyphis* chromosome 2, the supergene region (red lines), the region upstream (green lines), and the region downstream (blue lines) of the supergene. The yellow box marks a cluster of genes that appears to have been convergently fused to the social chromosome in both *Cataglyphis* and *Solenopsis* lineages (see Supplementary Figure S8).

Figure 5b plots the distributions of sizes of Ancestral Linkage Groups (ALGs) across chromosomes – groups of one-to-one orthologs that are conserved on the same chromosome across eight representative species of ants and bees. The general distribution of ALG sizes is roughly uniform across chromosomes, with inter-quartile ranges between 2 and 6, with few ALGs above 20 in some chromosomes. Notably, the largest ALG with 121 genes was found on the social chromosome, demonstrating the highest conservation of synteny in this chromosome. The 121 orthologous groups include 27 in the *Cataglyphis* supergene and 94 in the upstream region (blue in Figure 5c); 99 of the 121 are in the *Solenopsis* supergene, 5 upstream, and 17 downstream (Supplementary Table 2).

Figure 5c shows the conservation of microsynteny of genes on the social chromosome in three ants, two Apoidea species (the honey bee and a digger wasp), and the jewel wasp *Nasonia* as an outgroup (Supplementary Figure S8 shows all 22 Hymenoptera). Microsynteny in the supergene and the region upstream to the supergene in *Cataglyphis* appear to be conserved to a certain extent across the available genomes from Myrmicinae and Formicinae. In the clonal raider ant *Ooceraea biroi,* however, this region is broken up and represented on five chromosomes. The conservation of the chromosome in bees suggests that it was broken up in the lineage leading to *O. biroi* after it diverged from the lineage leading to Myrmicinae and Formicinae. It remains to be determined whether this is common to other members of the subfamily Dorylinae or specific to the clonal raider ant. The downstream region of the *Cataglyphis* chromosome is broken in many formicine species (e.g. *Formica*) and myrmicine species (e.g. *Pogonomyrmex*), and also in bees. Either the downstream region was part of the ancestral chromosome and broke off in multiple myrmicine and formicine lineages, or it was not part of the ancestral chromosome, and the same synteny group of genes was fused to the social chromosome in both *Cataglyphis* and *Solenopsis* lineages (marked in Figure 5c; see also Supplementary Figure S8).

Most genes in the upstream region of the *Cataglyphis* social chromosome and the supergene region (red and green lines in Figure 5c) are found on chromosome 15 of *Apis*, followed by chromosomes 1, 7, 8, and 14. The downstream *Cataglyphis* genes (blue lines) are not on *Apis* chromosome 15. This conservation of macrosynteny is evident across all the six representatives of the superfamily Apoidea, which includes the bees and the crabronid wasps (Supplementary Figure S8). These results indicate deep conservation of the social chromosome, dating back to the divergence of Apoidea and Formicidae (ants) around 175 million years ago^30^. This exceptional conservation of the social chromosome suggests that its genomic architecture has adaptive significance.

We propose that the conserved gene content in the supergene region is an ancestral genetic “toolkit” that has been repeatedly used in the evolution of sociobiological traits in hymenopteran lineages.

The association of social polymorphism with a supergene on the social chromosome in both *Cataglyphis* and *Solenopsis* can be interpreted in two ways. One possibility is that the supergene was present in the common ancestor and conserved in both lineages. However, the sequence divergence between the two *Solenopsis* supergene haplotypes —corresponding to only one million years ago— suggests a more recent origin^19,25^. Although the age of the *Cataglyphis niger* supergene remains unknown, no polygyne colonies have been reported from closely related *Cataglyphis* species (the bicolor group). This suggests that the two lineages independently evolved analogous supergenes on the same chromosome. While this may seem surprising, studies on sex chromosome turnover show that certain chromosomes repeatedly transition into sex chromosomes, likely due to the genes they carry^38,39^. Similarly, if the social chromosome harbors an ancient genetic toolkit for sociobiological traits, this would explain the repeated evolution of supergenes on this chromosome. Examining the genomic basis of multiple socially polymorphic species in other ant lineages could help shed light on this question.

### 2.5 Conserved gene content of the social chromosome across ants and bees

We examined the gene content of the *Cataglyphis* and *Solenopsis* social chromosomes to find 786 and 1088 putative protein-coding genes, respectively. The two supergene regions contain 248 and 441 putative protein-coding genes in *Cataglyphis* and *Solenopsis*, respectively. Next, we identified orthologous genes between the two chromosomes. In the *Solenopsis* supergene region, there are 207 functionally-annotated protein-coding genes with one-to-one orthologs in *Cataglyphis*, and 112 such genes in the *Cataglyphis* supergene, with 84 of these being found in both the supergene regions. All but one of the one-to-one orthologs of genes in the *Cataglyphis* supergene are in the *Solenopsis* social chromosome, and one gene is in an unassembled scaffold (Supplementary Table S3), demonstrating the exceptional degree of gene content conservation of this chromosomal region. Other genes have many-to-many orthology relationships between the two species (Supplementary Table S4; genes with functional annotation). The *Cataglyphis* social chromosome contains 492 such genes in total, of which 149 are in the supergene region. The *Solenopsis* social chromosome contains 665 such genes, 279 of which are in the supergene region.

The homologous supergene regions contain a large and diverse set of molecular functions. It is difficult to identify specific candidate genes without additional functional information. Nevertheless, we reviewed the list and looked for gene functions that may be implicated in the sociobiology of ants (Supplementary Tables S3 and S4 – one-to-one and many-to-many; Supplementary Figure S9 – dot plot with gene annotations).

One notable example is a cluster of three cytochrome P450 4C1 genes in the *Cataglyphis* supergene, which are orthologous to nine genes in the *Solenopsis* supergene. While the nine represent a statistically significant enrichment of P450 genes in the *Solenopsis* supergene (*p-*value = 0.013), the *Cataglyphis* supergene did not show significant enrichment (*p-*value = 0.13). Cytochrome P450 genes are a large and diverse family of enzymes, which was noted already when the first ant genomes were sequenced. The genome of the harvester ant *Pogonomyrmex barbatus* was found to contain 72 cytochrome P450 genes, which is a significant expansion relative to other insect species^40^. Enzymes of this gene family participate in a wide range of metabolic processes^41^, including synthesis and degradation of hormones such as juvenile hormone, a major regulator of insect development that plays a key role in caste determination in social insects^42–45^.

We also took note of the downstream region of the *Cataglyphis* chromosome, which is homologous to the upstream region (short arm) of the *Solenopsis* chromosome, because this region may have been convergently fused to the chromosome in both lineages (Figure 5c; Supplementary Figure S8). This region contains 121 functionally annotated one-to-one orthologous genes, including a steroid hormone receptor (*Cataglypis* gene_1712 / *Solenopsis* XP_039315095.1), an ecdysone 20- monooxygenase (*Cataglyphis* gene_2123 / *Solenopsis* XP_011172207.1), and an ecdysteroid-regulated 16 kDa protein (*Cataglyphis* gene_2111 / *Solenopsis* XP_011170763.1). Genes implicated in such functions may have significance for the sociobiology of ants, as different levels of ecdysteroids and ecdysone receptors were found in the brains of foragers versus nurses in *Camponotus* and *Harpegnathos*^46,47^.

The total number of functionally annotated one-to-one orthologs shared between the *Cataglyphis* and *Solenopsis* social chromosomes and *Apis mellifera* (anywhere in the bee genome) is 305. Of these, 164 (54%) are located on *Apis* chromosome 15 (Supplementary Table S5), and 34 out of the 164 are located in the supergenes of both *Solenopsis* and *Cataglyphis*. These 34 genes include dopamine receptor 2 (*Cataglypis* gene_1588 / *Solenopsis* XP_039314227.1 / *Apis* XP_026301043.1), odorant binding protein-9 (*Cataglyphis* gene_1588 / *Solenopsis* XP_039314227.1 / *Apis* XP_026301043.1), and neuronal acetylcholine receptor subunit alpha-10 (*Cataglyphis* gene_2000 / *Solenopsis* XP_025994007.1 / *Apis* XP_392070.3). Dopamine is implicated in modulating social and reproductive behavior in Hymenoptera^48–52^. List of many-to-many orthologs between the three species are in Supplementary Table 6.

### 2.6 Enrichment in olfactory and neural signaling functions in the social chromosome across ants and bees

We tested for enrichment of 13,317 Gene Ontology (GO) functional categories in the conserved gene set on the social chromosome. We analyzed representative genomes from eight distantly related Formicidae and Apoidea lineages (ants and bees). We defined a conserved set of 187 orthologous gene groups (OGs) where the ortholog is found in the chromosome homologous to the social chromosome in at least six out of the eight genomes. Table 1 lists the GO categories that were significantly enriched (FDR corrected *p*-value < 0.05). The full table with additional information can be found in Supplementary Table S7. Most notably, the top GO categories include olfactory learning and response to odorants (GO: 0090328, GO: 1990834). Other enriched GO categories are related to neural functions, especially dopamine signaling (GO:0007191, GO:0004952, GO:0001588). These results indicate that the conserved gene set in the social chromosome is involved in chemical communication and neural signaling. Other GO categories that were significantly enriched were involved in metabolic regulation and immune function. There was no significant GO enrichment in the set of 83 OGs in the overlapping region of the *Cataglyphis* and *Solenopsis* supergenes. The conservation of the social chromosome across ants and bees suggests that the homologous chromosome may also code for sociobiological functions in bees. To begin to investigate this, we used the set of genes that were inferred to be under positive selection in bee lineages where higher levels of sociality evolved ^53^. However, a test for enrichment of this gene set in *Apis* chromosome 15 did not detect enrichment. While this analysis gave a negative result, there are many other datasets of genes related to bee sociobiology that could be investigated in future studies.

**Table 1:** Functional categories enriched in the conserved gene set of the social chromosome

| GO category | OGs with GO* | FDR corrected p-value | *Cataglyphis* gene in each conserved OG |
| --- | --- | --- | --- |
| BP: Regulation of olfactory learning (GO:0090328) | 3/5 | 0.019 | Dopamine receptor 1 (gene_1435), Octopamine receptor in mushroom bodies *Oamb* (gene_1830), Uncharacterized protein (gene_1662) |
| BP: Response to odorant (GO:1990834) | 3/7 | 0.019 | Dopamine receptor 1 (gene_1435), Dopamine receptor 2 (gene_1588), Uncharacterized protein (gene_1662) |
| BP: Adenylate cyclase-activating dopamine receptor signaling pathway (GO:0007191) | 2/2 | 0.019 | Dopamine receptor 1 (gene_1435), Dopamine receptor 2 (gene_1588) |
| MF: Dopamine neurotransmitter receptor activity coupled via Gs (GO:0001588) | 2/2 | 0.019 | Dopamine receptor 1 (gene_1435), Dopamine receptor 2 (gene_1588) |
| MF: Dopamine neurotransmitter receptor activity (GO:0004952) | 2/2 | 0.019 | Dopamine receptor 1 (gene_1435), Dopamine receptor 2 (gene_1588) |
| BP: Positive regulation of sensory perception of pain (GO:1904058) | 2/2 | 0.019 | Tachykinin-like peptides receptor 99D (gene_1948), ATP synthase subunit C lysine N-methyltransferase (gene_1859) |
| BP: Propionate metabolic process methylcitrate cycle (GO:0019679) | 2/2 | 0.019 | 4-hydroxybutyrate coenzyme A transferase (gene_1884, gene_1885) |
| BP: Acetate metabolic process (GO:0006083) | 2/2 | 0.019 | 4-hydroxybutyrate coenzyme A transferase (gene_1884, gene_1885) |
| MF: Acetate CoA-transferase activity (GO:0008775) | 2/2 | 0.019 | 4-hydroxybutyrate coenzyme A transferase (gene_1884, gene_1885) |
| BP: Positive regulation of activated T cell proliferation (GO:0042104) | 2/2 | 0.019 | Signal transducer and activator of transcription 5B (gene_1887), Glycerol-3-phosphate acyltransferase 1 mitochondrial (gene_2005) |
| BP: Positive regulation of programmed cell death (GO:0043068) | 2/2 | 0.019 | Cell death specification protein 2 (gene_1943), Uncharacterized protein (gene_1662) |
| CC: Integral component of membrane (GO:0016021) | 33/1241 | 0.019 | 33 genes including Dopamine receptor 1 (gene_1435), Dopamine receptor 2 (gene_1588), Octopamine receptor in mushroom bodies *Oamb* (gene_1830), Tachykinin-like peptides receptor 99D (gene_1948), ATP synthase subunit C lysine N-methyltransferase (gene_1859), Serine protease HTRA2 mitochondrial (gene_1443), Attractin-like protein 1 (gene_1831), Pickpocket protein 28 (gene_1847), Beta-14-galactosyltransferase 7 (gene_1850), ABC transporter G family member 20 (gene_1862), MAP kinase-activating death domain protein (gene_1873), probable sodium/potassium/calcium exchanger CG1090 (gene_1906), Calcium-independent phospholipase A2-gamma (gene_1530), Sphingosine-1-phosphate lyase (gene_1949), Tetraspanin-33 (gene_2003), Sodium-coupled neutral amino acid transporter 9 (gene_1480) |
| MF: Manganese ion binding (GO:0030145) | 4/24 | 0.027 | Beta-14-galactosyltransferase 7 (gene_1850), m7GpppN-mRNA hydrolase (gene_1532), NADP-dependent malic enzyme (gene_1951), DNA-directed primase/polymerase protein (gene_1937) |
| MF: Hydrolase activity (GO:0016787) | 5/49 | 0.041 | V-type proton ATPase subunit B (gene_1502), DNA repair and recombination protein RAD54-like (gene_1503), Fumarylacetoacetate hydrolase domain-containing protein 2A (gene_2029), Phosphoglycerate mutase 2 (gene_1893), Uncharacterized protein (gene_2004) |
| MF: Acetyl-CoA hydrolase activity (GO:0003986) | 2/3 | 0.041 | 4-hydroxybutyrate coenzyme A transferase (gene_1884, gene_1885) |
| BP: Regulation of presynaptic cytosolic calcium ion concentration (GO:0099509) | 2/3 | 0.041 | Dopamine receptor 1 (gene_1435), Dopamine receptor 2 (gene_1588) |

Results of the hypergeometric test for enrichment of Gene Ontology (GO) categories in the conserved set of Ortholog Groups (OGs) that are on the social chromosome in at least six out of the eight species included in the analysis.
BP = biological process; MF = molecular function; CC = cellular component.
\* The number of OGs in the conserved gene set out of the total number of OGs in the dataset that were annotated with a specific GO term.

We were particularly interested in genes implicated in olfactory functions, which were extensively studied in *Solenopsis* because of their potential role in chemical signaling and the regulation of social structure. Previous studies reported differences between monogyne and polygyne queens in cuticular hydrocarbons and other compounds that were suggested to function as queen pheromones involved in the regulation of social form and the green beard effect^35,54,55^. Other studies focused on odorant binding proteins (OBPs) and odorant receptors (ORs) in the *Solenopsis* supergene, which may be involved in olfactory response to such chemical signals^56–59^. The *Solenopsis* supergene contains a large array of 23 ORs and 3 more separate ORs^59^. We annotated ORs in the *Cataglyphis niger* genome and constructed a gene tree including ORs from *C. niger, Solenopsis invicta, Camponotus floridanus,* and *Linepithema humile* (Supplementary Figure S9). The 23 *Solenopsis* ORs and their orthologs in *Cataglyphis* belong to a single subfamily of the OR gene superfamily. The gene tree shows that the 23 *Solenopsis* ORs have one-to-one or many-to-many orthology relationships with ORs in two regions of the *Cataglyphis* social chromosome, but outside of the supergene (marked by red stars on the dot plot in Figure 4c). Three more ORs in the *Solenopsis* supergene (belonging to a different OR subfamily) are orthologs of two *Cataglyphis* ORs, which also reside outside of the supergene. Given that the ORs are a fast-evolving family, there is no clear orthology relationship between ant and bee ORs.

The *Solenopsis* supergene also contains 9 out of a total of 23 OBPs in the genome^58^. One of the OBPs is *Gp-9*, the first gene discovered in the *Solenopsis* supergene in the 1990s and the one protein that shows the largest number of amino acid substitutions between the M and the P chromosomes, indicating positive selection^56–58^. Another OBP in the *Solenopsis* supergene was duplicated and shows polygyne-specific expression^58,60^. Interestingly, the homologous chromosome in the honey bee, chromosome 15, also contains 9 out of 21 OBPs, but only one of these (AmelOBP9) is orthologous to one of the supergene OBPs (SiOBP9). The 9 OBPs in the *Solenopsis* supergene are orthologous to 3 OBPs in the *Cataglyphis* supergene (Fig. 4c).

Cnig_gene_1607 is orthologous to 7 of the *Solenopsis* OBPs; Cnig_gene_1608 is the ortholog of the *Solenopsis Gp-9* gene (SiOBP3); and Cnig_gene_1586 is orthologous to SiOBP9 and AmelOBP9 (Fig. 6). We note that the OBP cluster containing *Gp-9* is in the largest block of conserved microsynteny between *Solenopsis* and *Cataglyphis*. The 3, 9, and 9 OBPs in the *Cataglyphis*, *Solenopsis*, and *Apis* chromosomes represent a statistically significant enrichment (p-value = 0.0017, 9.28e-9, and 1.17e-06, respectively). Altogether, these gene families implicated in olfaction are promising candidates for the toolkit that may have been reused for social evolution in multiple hymenopteran lineages.

**Figure 6:**
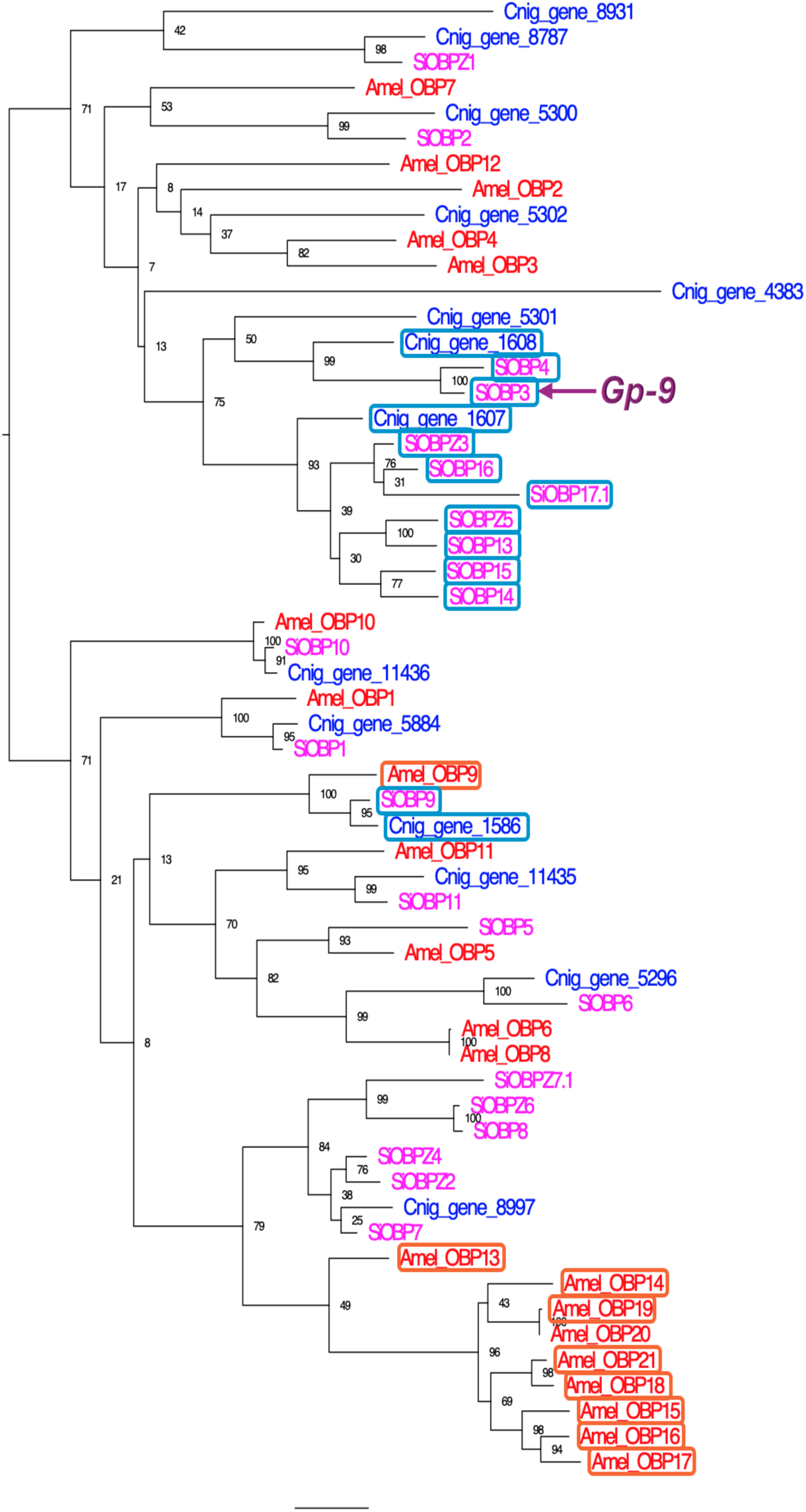
Gene tree of the odorant binding protein (OBP) gene family in the ants *Cataglyphis niger* and *Solenopsis invicta* and the honey bee *Apis mellifera*. OBPs marked by rectangles reside on the social chromosomes (*Cataglyphis* chromosome 2, *Solenopsis* chromosome 16, *Apis* chromosome 15).

## 3. Conclusion

The complex trait of multi-queen (polygyne) colonies has evolved repeatedly from single-queen (monogyne) colonies in ants^8,9^. Multiple ant species display polymorphism in social form, providing a unique opportunity to investigate the evolutionary genomic basis of this trait. We investigated the genomic architecture associated with social form in a polymorphic population of *Cataglyphis niger*. Our data show that social polymorphism is associated with a large supergene— a region of suppressed recombination across at least 5.8Mb of chromosome 2. This supergene is virtually always heterozygous (MP) in polygyne individuals and homozygous (MM) in monogynes. Surprisingly, this supergene is homologous to the supergene of the fire ant *Solenopsis invicta*, even though these two lineages diverged more than 90 million years ago. These species are representatives of the two largest ant subfamilies Formicinae and Myrmicinae. The low sequence divergence between supergene haplotypes suggests that they evolved relatively recently in these two distant lineages, implying that this is a case of convergent evolution. Most of *Cataglyphis* chromosome 2 is homologous to most of *Solenopsis* chromosome 16, with a substantial overlap of the supergene regions where the gene order is highly conserved. The conservation of this set of genes suggests that this chromosome encodes functions related to social colony structure and the sociobiology of ants that is likely to be shared across many ant species. Conservation of synteny in gene content of the social chromosome is also observed in representatives of Apoidea, including the honeybee *Apis mellifera*. This chromosome contains the largest Ancestral Linkage Group (ALG) across the genome, consisting of 121 genes with conserved linkage across ants and bees. Olfactory functions are overrepresented in this conserved gene set. We highlight the large clusters of odorant binding proteins (OBPs) in the homologous chromosomes of *Apis* honeybees and *Solenopsis* fire ants. These genes were implicated by evolutionary and gene expression studies in the social polymorphism in *Solenopsis*. The observation of conserved synteny dating back to around 175 MYA indicates that this set of genes acted in synergy to produce some traits with fitness consequences. Specifically in ants, this gene set facilitated the repeated evolution of polygyne social structure. The conservation of synteny before the evolution of social polymorphism in *Solenopsis* and *Cataglyphis* suggests that this gene set was responsible for some traits of consequence over 175 million years, possibly also related to sociobiology. The conserved gene set could have served as a genetic “toolkit” for evolving sociobiological traits across bee and ant lineages. This discovery opens the way for both evolutionary and mechanistic studies of the genetic basis and genomic architecture of complex phenotypes related to insect sociality.

## 4. Methods

### 4.1 Sample collection

Thirty nests of *Cataglyphis niger* were collected from a single population in Tel Baruch beach, Israel, between November and December 2019 (Supplementary Table 1). This location is known to harbor both monogyne and polygyne colonies based on Reiner Brodetzki *et al.*^32^. We targeted at least two monogyne and two polygyne nests in every sampling trip based on their geographical distribution in that previous study. This also allowed us to conduct aggression assays (see below) between monogyne and polygyne nests after every batch of sampling. A distance of at least 100m was maintained between two sampled nests as these ants are known to be polydomous. An attempt was made to collect the queens by excavating the entire nest. Nest samples were carried back to the lab (TBA14 - 44) to perform behavioral experiments and extract genomic DNA. An additional polygyne nest (TBA67) with several queens was collected in January 2021 and was used for building a linkage map. The queens from this nest were isolated and 225 samples from the brood of a single queen were collected for RAD sequencing.

### 4.2 Molecular work

Samples not used for the behavioral assay were stored at -80C immediately after sampling. 22-24 workers from each nest in addition to the queens amounting to 714 samples were used for the molecular work from sampling carried out in 2019, henceforth referred to as the SC (social chromosome) dataset. An additional 225 brood samples from a single polygyne mother were used to construct a linkage map (linkage map dataset). DNA was extracted from the workers’ abdomen and the queen’s thorax using a liquid handling robot (Tecan Freedom EVO 150 NGS) and DNeasy (QIAGEN) protocol. The extracts were cleaned using ethanol precipitation and stored at -20C. A partial fragment of 450bp of Cytochrome b (*cyt b*) was PCR amplified and Sanger sequenced for 2-3 workers and queens from each nest. Double-digest Restriction-site Associated DNA sequencing (RAD-seq) was used to generate genomic libraries following Brelsford *et al.*^61^. Briefly, the genomic DNA was cut using restriction enzymes EcoRI and BfaI. Adaptors with individual barcodes were ligated to each sample for multiplexing. The ligated products were PCR amplified using barcoded primers and pooled. Size selection of DNA fragments between 330–1000bp was carried out using a Bluepippin instrument (Sage Science). A detailed protocol can be found in Lajmi *et al.*^62^. Libraries were sequenced on an Illumina HiSeq X machine with paired-end 150bp reads.

### 4.3 Molecular Analysis

#### 4.3.1 Mitochondrial phylogeny

The Sanger sequences were aligned in MEGA X^63^ using the Muscle algorithm^64^. To assign our samples to previously described mitotype clades, published *cyt b* sequences of monogyne and polygyne colonies were also included from Reiner Brodetzki *et al.*^32^. Mitotypes of *Cataglyphis israelensis* were used as an outgroup. A maximum likelihood tree was constructed using raxmlGUI 2.0^65^ using a GTR+G model of sequence evolution and 1000 bootstraps.

#### 4.3.2 RAD-seq pipeline

The bioinformatics pipeline was run separately for the SC data and linkage map, as well as the two datasets combined. The raw sequencing data were examined using FastQC^66^ for quality. Adaptors and low-quality reads were then trimmed/filtered using Trimmomatic^67^. Two mismatches were allowed in identifying the adaptors, with eight being the minimum length for identifying an adaptor, a sliding window size of 4bp with an average required quality of 15 was used. FLASH^68^ was used to combine overlapping paired reads. The data were then demultiplexed using the command process_radtags.pl from the Stacks2 package^69^. The reads were then mapped to the *Cataglyphis niger* genome (NCBI GenBank assembly GCA_050437125.1 ^70^) using Bowtie2^71^ with the -- end-to-end and --very-sensitive options. SNP calling was conducted using Ref_map.pl from Stacks2 to obtain a VCF file. Samples with more than 50% missing data were dropped from the analyses. The SNP calling was done separately for the SC and linkage map datasets, as well as the combined dataset. A blacklist of loci with potential artifacts was created based on the heterozygous genotype calls in three or more samples out of 64 haploid males. These loci were excluded from the VCF. Such artifacts may be the result, for example, of errors in the assembly of the reference genome, where repetitive sequences are collapsed into a single copy.

Along with removing the blacklisted loci, the following loci were removed from the VCF file for the SC dataset using VCFtools version 0.1.14^72^: i) genotype calls based on less than 12 reads; ii) loci with more than 30% missing data; iii) loci with a minor allele frequency of less than 0.4% or 3 allele copies; iv) loci with a mean depth greater than 175 reads per individual. The final VCF file for the SC dataset consisted of 672 individuals comprising 28 queens and 644 workers. This resulted in a catalog of 30,955 high-confidence SNPs that were used for most of our analyses. The linkage map data VCF file varied slightly in the filtration in that we required only 8 reads for a genotype call, a maximum mean depth of 150 reads per individual (because of lower average depth), and a minor allele frequency of 1% or 3 allele copies (because of the smaller number of samples). The final VCF file for the linkage map consisted of 225 offspring and their mother and amounted to 17,984 high-confidence SNPs.

### 4.4 Identification of the social form

*Cataglyphis niger* is a polydomous species, i.e., a colony may have multiple nests. This made finding the queens of each colony difficult. Therefore, we relied on kinship estimates among worker samples to identify the colony’s social form. Monogyne nests were expected to show high kinship between nestmates as they are expected to be full or half-siblings, compared to polygyne nests where workers are not necessarily related. We also conducted behavioral assays, as additional supporting evidence for the colony’s social structure.

#### 4.4.1 Kinship

Kinship inference was carried out for the two genomic datasets separately. For the SC dataset, kinship was used to identify the social form of the nests. Whereas in the linkage map dataset, kinship was used to identify the patrilines. The RAD-seq SNP catalog was imputed and phased using Beagle version 5.0^73^ and refined-IBD^74^ was used to identify shared identity-by-descent (IBD) haplotypes between samples. The kinship between a pair of samples was estimated based on the combined lengths of their shared IBD haplotypes, resulting in a kinship matrix. To obtain an estimate of the relatedness coefficient we divided the total IBD length by twice the total genome assembly length (because the samples are diploid). A total of 7,651,126 and 6,238,277 IBD haplotypes were identified in the SC dataset (among the 672 samples) and the linkage map dataset (among the 225 samples), respectively. Social form was assigned to each nest based on the distribution of kinship scores. Nests were classified as polygyne if more than 75% of pairs of nestmates had less than 20% relatedness, and as monogyne otherwise. To ensure that the high relatedness in monogyne nests were not due to higher inbreeding, we calculated individual FIS for one sample per colony. We found that individual inbreeding coefficients did not differ significantly between monogyne (median FIS = 0.073, IQR = 0.068–0.089, n = 13) and polygyne nests (median FIS = 0.070; IQR = 0.060–0.110, n = 17; Wilcoxon rank-sum test, W = 120, p-value = 0.711), indicating that both the social forms are largely outbred. We also checked the kinship between workers and queens in the five nests that were assigned monogyne social form based on kinship results and where only one queen was found. As expected, we found found that relatedness values were very close to 50% (Supplementary Figure S1).

#### 4.4.2 Behavioral experiment

Behavioral assays were conducted within a week of collection from the field. These assays were based on the assumption that workers from different colonies would show high aggression towards each other. Therefore, workers from two different monogyne nests were expected to show higher aggression than workers from two polygyne nests, as we expect all polygyne ants to be part of a supercolony^32^. Dyadic aggression assays were performed, where two ants from different nests (test) or the same nest (control) were introduced into a round 12 cm diameter arena and their behavior was video recorded for three minutes. Behavior aggression scores were coded as antenation=0, mandibular threat=1, short bite with jump=2, bite=3, spraying=4 and the total aggression score was calculated as

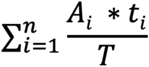

where Ai and ti are the aggression score and duration of each behavior over the total time T, respectively. Ten trials were conducted for each pair of nests that was tested and an average score was calculated. A total of 1029 assays were carried out to get 104 averaged scores between pairs of nests. A two-sample *t*-test was conducted to compare the aggression scores between nestmates, polygyne-polygyne nests, and monogyne/polygyne-monogyne nest. The test was performed in R using command t.test.

### 4.5 Identifying genomic regions associated with social form

Weir and Cockerham (1984)^75^ *FST* was calculated for every SNP using VCFtools version 0.1.14^72^ between 282 monogyne samples and 388 polygyne samples, where the social form was defined based on the distribution of kinship score within the nest. Linkage disequilibrium (LD; *r*^2^) analysis was performed among SNPs in each chromosome for monogyne and polygyne samples separately, using Plink v1.9^76^. Only SNPs exhibiting a minor allele frequency surpassing 0.05 for each social form were included in this analysis.

#### 4.5.1 Linkage analysis in the progeny of a polygyne queen

To test for recombination in the supergene in polygyne colonies, we isolated a single polygyne queen from nest TBA67 with workers (TBA67c). 269 samples consisting of larvae and pupae were collected in a span of 60 days after the queen was isolated. RAD-seq libraries were constructed for the queen and her offspring using the same protocol as above. Patrilines were identified using kinship analysis (as above) and a linkage map was constructed for the patriline with the largest number of samples (69 samples). A linkage map was constructed based on 3,149 bi-allelic loci where the minor allele was present in at least two individuals from the same patriline. However, this analysis failed in the supergene region of chromosome 2 because all 69 samples had the same genotype in every SNP. Therefore, it was not possible to estimate the frequency of recombination and test for linkage between neighboring SNPs. To illustrate this unusual genetic pattern, we plotted the heterozygosity in the 69 progeny along the chromosome for all SNPs that were heterozygous in the mother queen.

### 4.6 Synteny analysis

Synteny analysis was carried out to evaluate the conservation of synteny in the chromosome carrying the supergene across 22 hymenopteran species with the oldest divergence dating to ∼230 million years ago. The analysis included the ant subfamilies Formicinae (*Cataglyphis niger*, *C. hispanica*, a *Formica aquilonia x polyctena* hybrid*, Camponotus floridanus,* and *Lasius platythorax*), Myrmicinae (*Monomorium pharaonis*, *Solenopsis invicta*, *Pogonomyrmex californicus,* and *Trachymyrmex septentrionalis*) and Dorylinae (*Ooceraea biroi*). Six species belonging to the Apoidea superfamily, the sister clade to ants, were also included in this analysis. This included bee families Apidae (*Apis mellifera*) and Colletidae (*Colletes gigas*), a solitary bee belonging to Megachilidae (*Osmia bicornis*), and solitary wasps belonging to Crabronidae (*Pemphredon lugubris*, *Nysson spinosus*, and *Ectemnius continuus*). Six additional wasp lineages were included— families Vespidae (*Vespula vulgaris* and *Ancistrocerus nigricornis*), Chalcidoidea (*Nasonia vitripennis*), Ichneumonidae (*Venturia canescens*), Ichneumonoidea (*Mocroplitis mediator*), and Pompilidae (*Anoplius nigerrimus)*. The genomes for this analysis were chosen based on their availability on NCBI, a scaffold N50 of more than 1MB, and gene annotation for the genome assembly (Supplementary table 9). OMA v2.6.0^77^ was used to infer orthogroups and transfer functional annotations from *Solenopsis invicta* orthologs to *Cataglyphis niger* genes.

We also used the ODP algorithm^78^ to identify Ancestral Linkage Groups (ALGs) with conserved synteny in chromosome-level assemblies of eight representatives of divergent Formicidae and Apoidea lineages: *Ectemnius continuus*, *Pemphredon lugubris*, *Colletes gigas*, *Apis mellifera*, *Pogonomyrmex californicus*, *Solenopsis invicta*, *Cataglyphis niger*, *Lasius platythorax*.

### 4.7 GO enrichment analysis

GO categories were assigned to *C. niger* genes using Trinotate^79^ and used to test for enrichment on the social chromosome using the hypergeometric test implemented in the python module scipy^80^. We used Hierarchical Orthologous Groups (HOGs) defined by OMA for the same chromosome-level assemblies of eight representatives of Formicidae and Apoidea as in the ODP analysis above. From the OMA results, HOGs uniqely presented on at least on six out of the eight genomes in the chromosome homologous to *C. niger* chromosome 2 were designated as our sample population for the test. For each GO category present on these HOGs, we counted how often they are present in these compared to the whole genome.

### 4.8 OR and OBP phylogenetic analysis

Odorant receptors (ORs) were annotated using HAPpy-ABCENTH^81^, based on manually curated OR gene models from 5 ant species. The first part of the program locates putative genetic loci through homology, and the second part, ABCENTH, is a gene finder specifically built for multigene families like ORs, with high sequence divergence but a very conserved exon structure. OR protein sequences were compiled from *Cataglyphis niger, Solenopsis invicta*, *Linepithema humile,* and *Camponotus floridanus.* Odorant Binding Protein (OBP) sequences for *Apis mellifera*^82^ and *Solenopsis invicta*^58^ were downloaded from NCBI and *Cataglyphis* annotations were used from Inbar *et al.*^70^ and analyzed separately. For each of these datasets, MAFFT^83^ was used to build a multiple sequence alignment. OR and OBP gene trees were reconstructed using RAxML v8.2.12^84^, with the PROTCALG model, the LG rate matrix, and 100 bootstraps repeats.

## Acknowledgments

We thank Abraham Hefetz for introducing us to the polymorphic study population and for background knowledge on this species and its social forms. We are grateful to Jessica Purcell and Alan Brelsford for their advice on RAD sequencing and the study of supergenes. Brian Charlesworth, Gene Robinson, Laurent Keller, and Lukas Schrader, for their insightful and constructive comments on the manuscript. EP would like to thank Laurent Keller for inspiring and motivating the study of social supergenes. AL would like to thank Praveen Karanth for his support during the analysis and writing of this paper.

AL was supported by the “Study in Israel” Fellowship provided by the Planning & Budgeting Committee (PBC) of the Council for Higher Education (CHE). This study was funded by Israel Science Foundation grants 646/15 and 1845/22.

## 6. Supplementary material

**Supplementary Figure S1:** (a) Relatedness scores within five monogyne nests where queens were present. Mother-daughter pairs (wQ) have relatedness of around 50% whereas worker-worker pairs (ww) have around 25% and 75% relatedness in most cases. (b) relatedness scores within nests identified as monogyne and polygyne. (c) Distribution of relatedness scores within each nest.

**Supplementary Figure S2:**
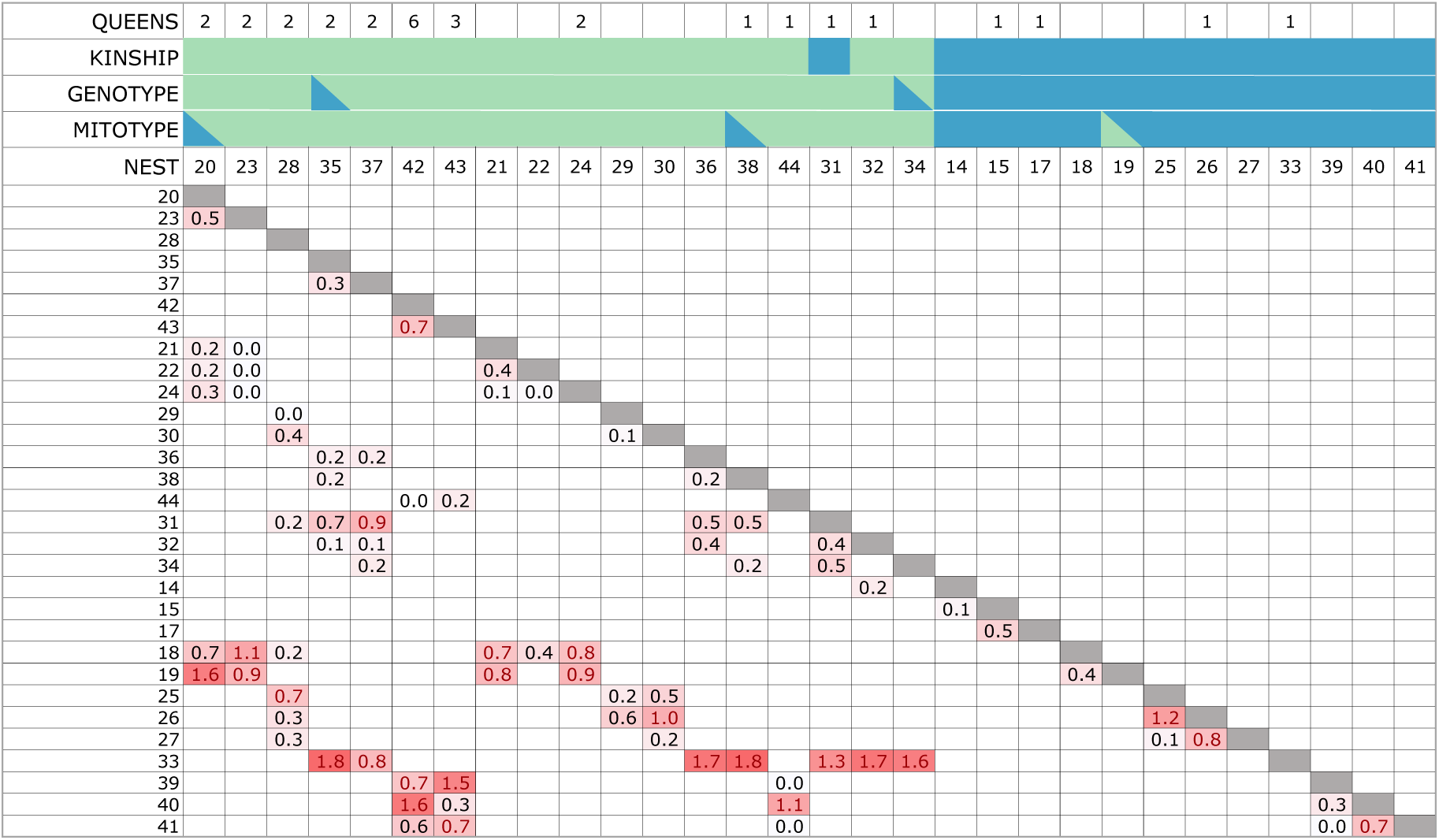
Matrix of average aggression scores among nests. The intensity of cell color is proportional to the aggression score. The first four rows provide nest IDs, kinship, genotype at the supergene, number of queens found, and mitotypes (monogyne/polygyne). The monogyne form is denoted in blue and polygyne in green.

**Supplementary Figure S3:**
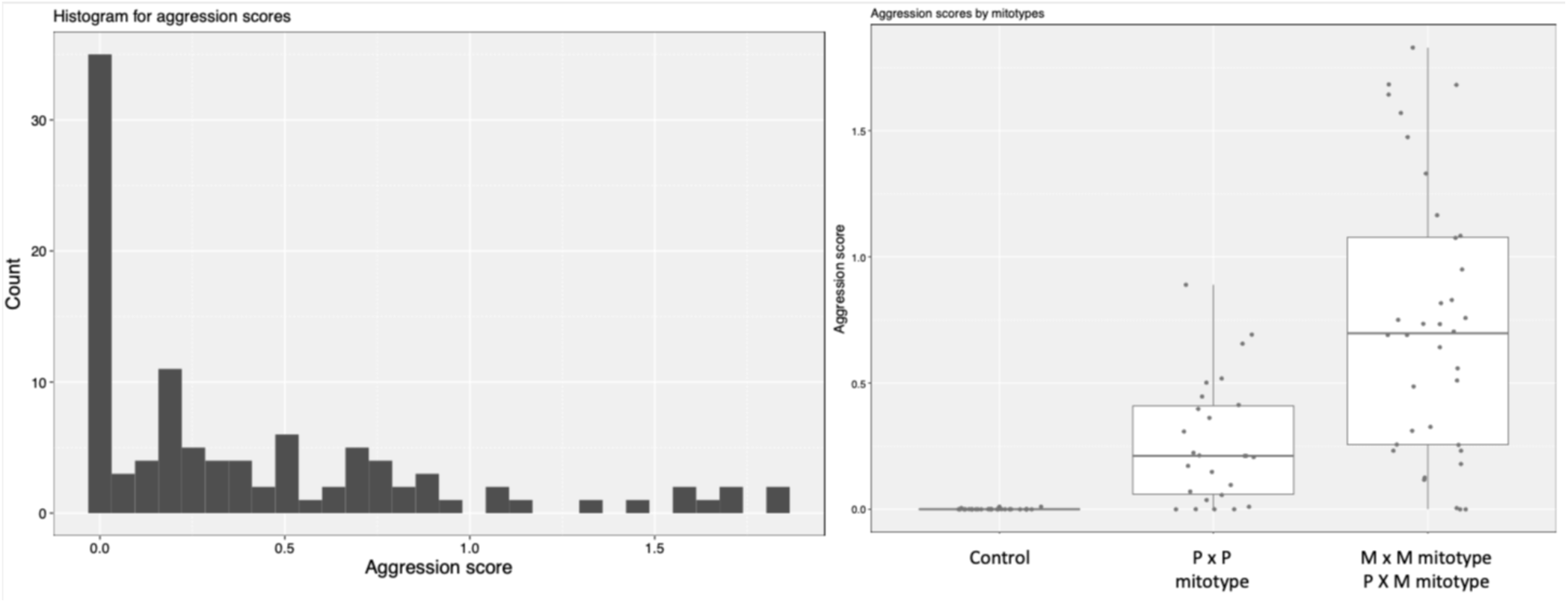
(a) Histogram of aggression scores. (b) Aggression scores across social forms assigned based on mitotype.

**Supplementary Figure S4:**
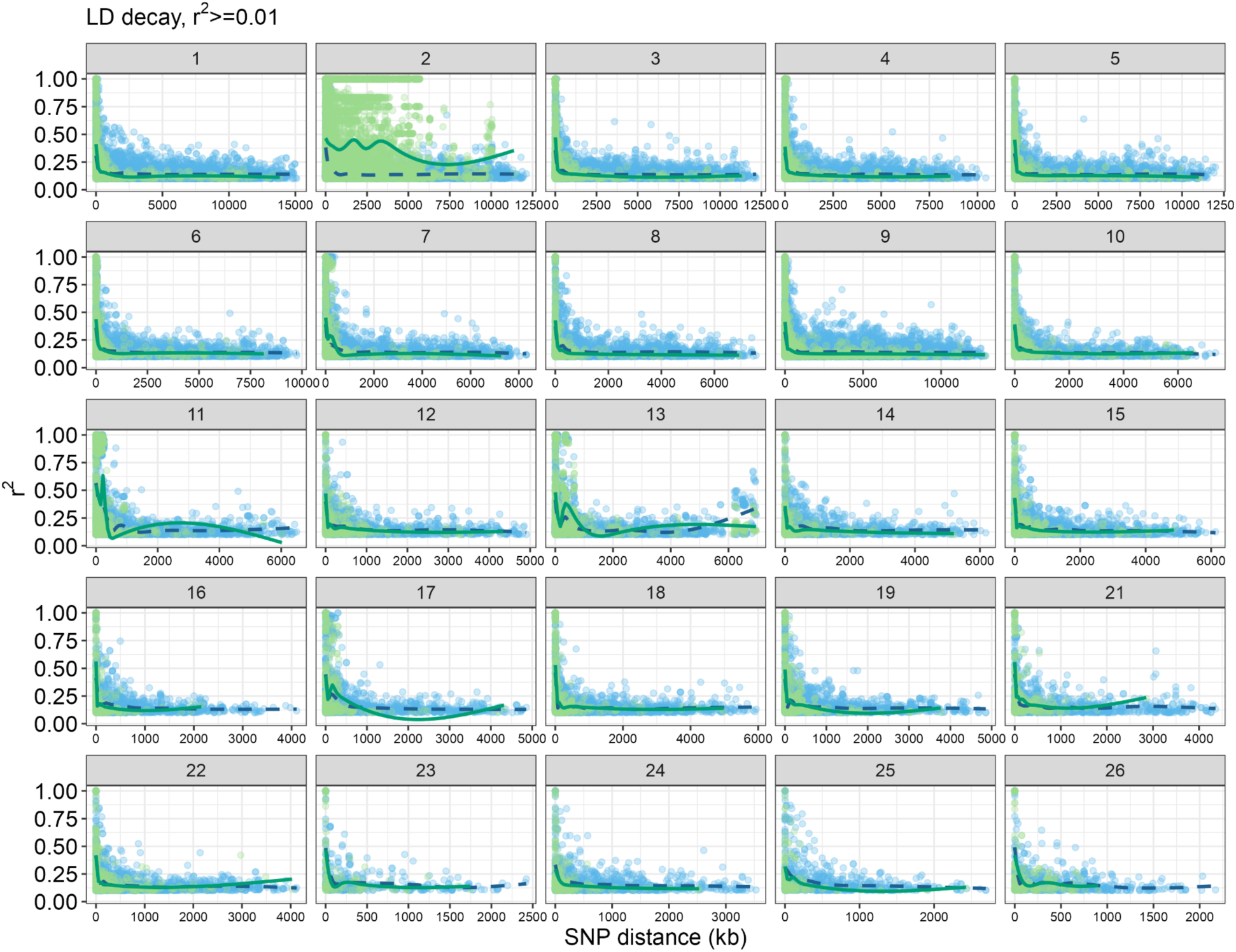
LD decay plot for *Cataglyphis niger* chromosomes. Blue dots represent the LD at corresponding SNP distances in monogyne samples, while green dots represent polygyne samples. The blue dashed and solid green lines indicate the average LD at each SNP distance in monogyne and polygyne samples, respectively.

**Supplementary Figure S5:**
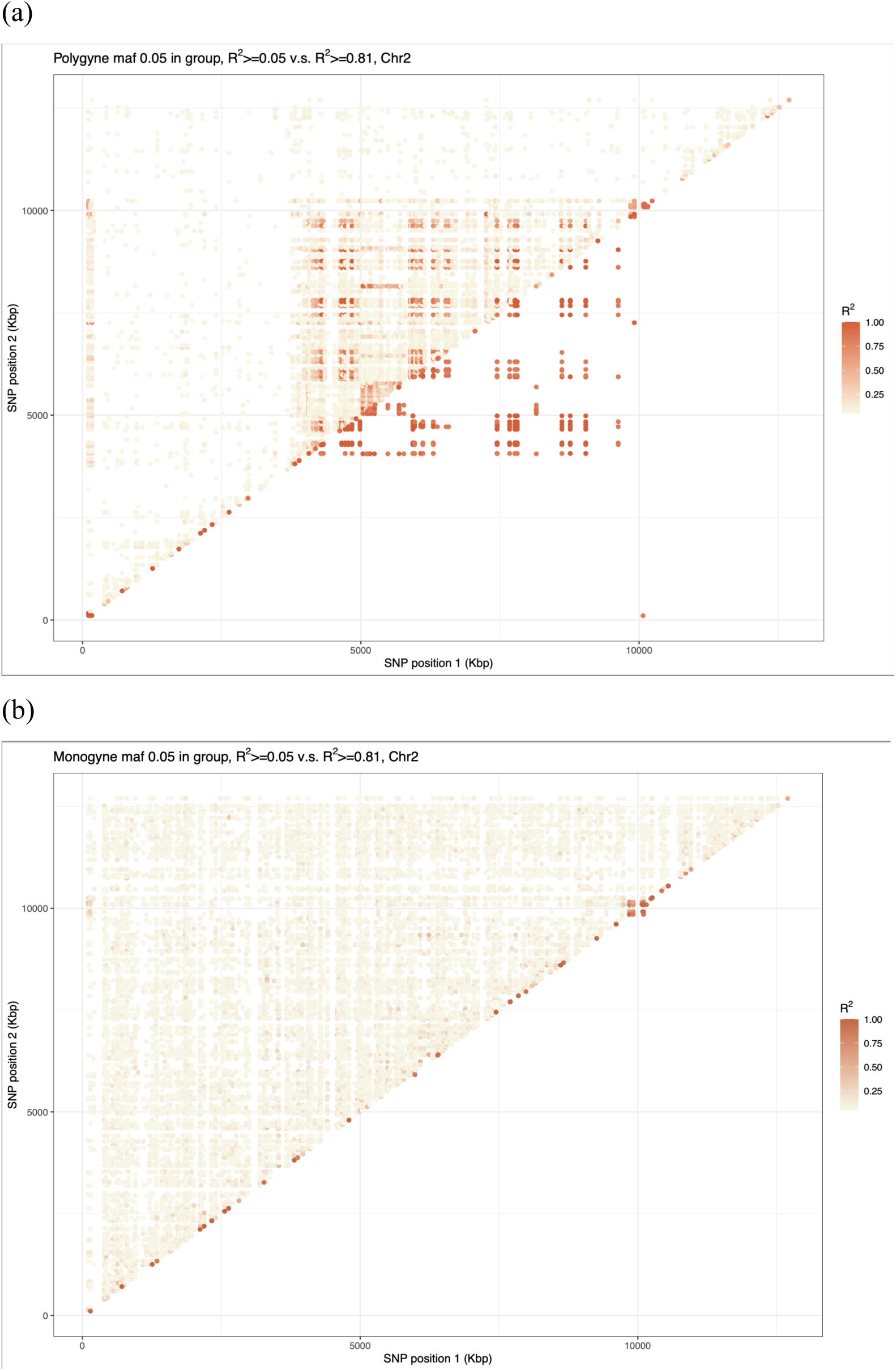
A plot of linkage disequilibrium (LD) along chromosome 2 in (a) polygyne and (b) monogyne samples.

**Supplementary Figure S6:**
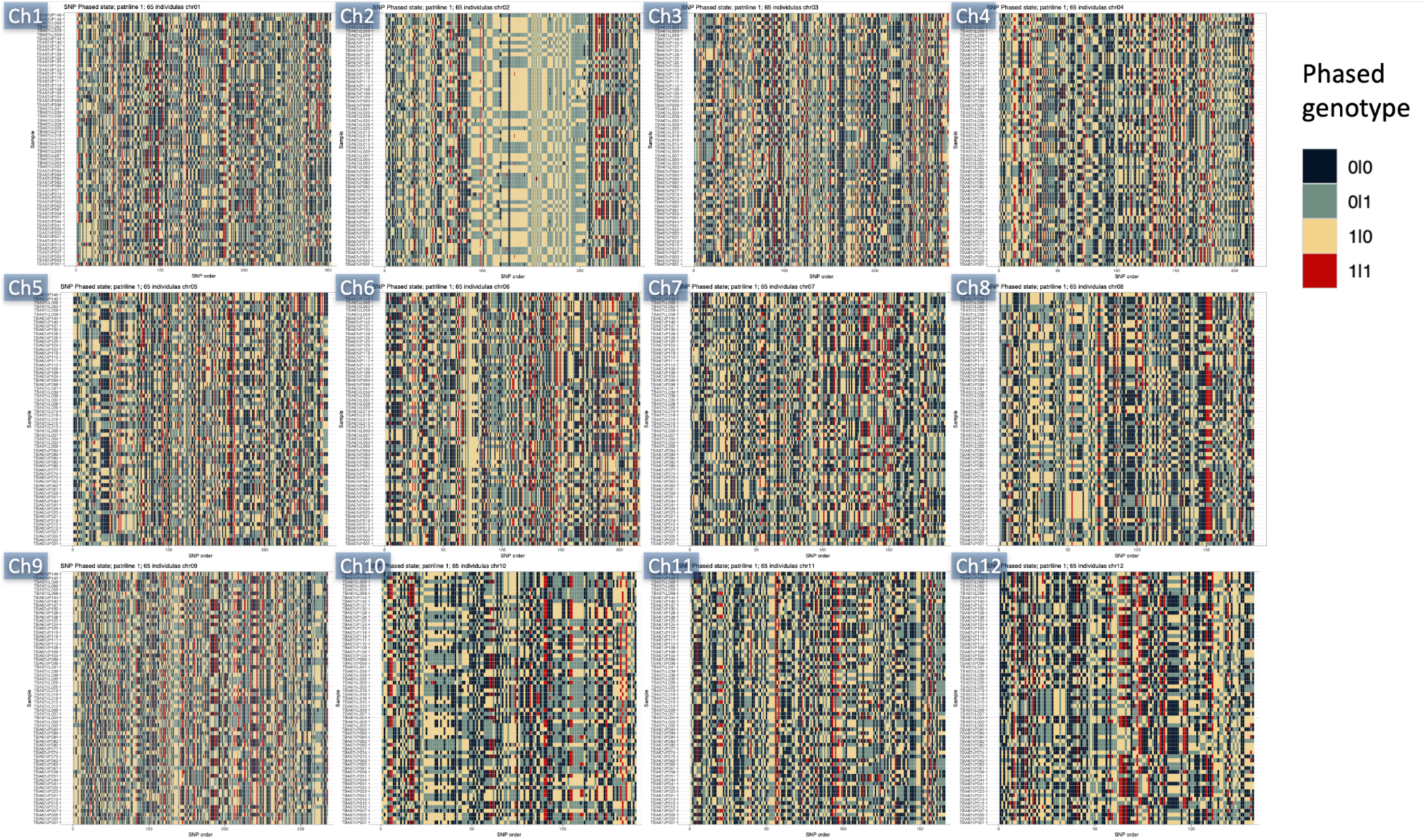
Phased genotypes of 65 daughters of a single polygyne queen and a single father (full-sisters). Only sites that were polymorphic in these samples are shown. The supergene region in chromosome 2 stands out for having 100% heterozygosity.

**Supplementary Figure S7:**
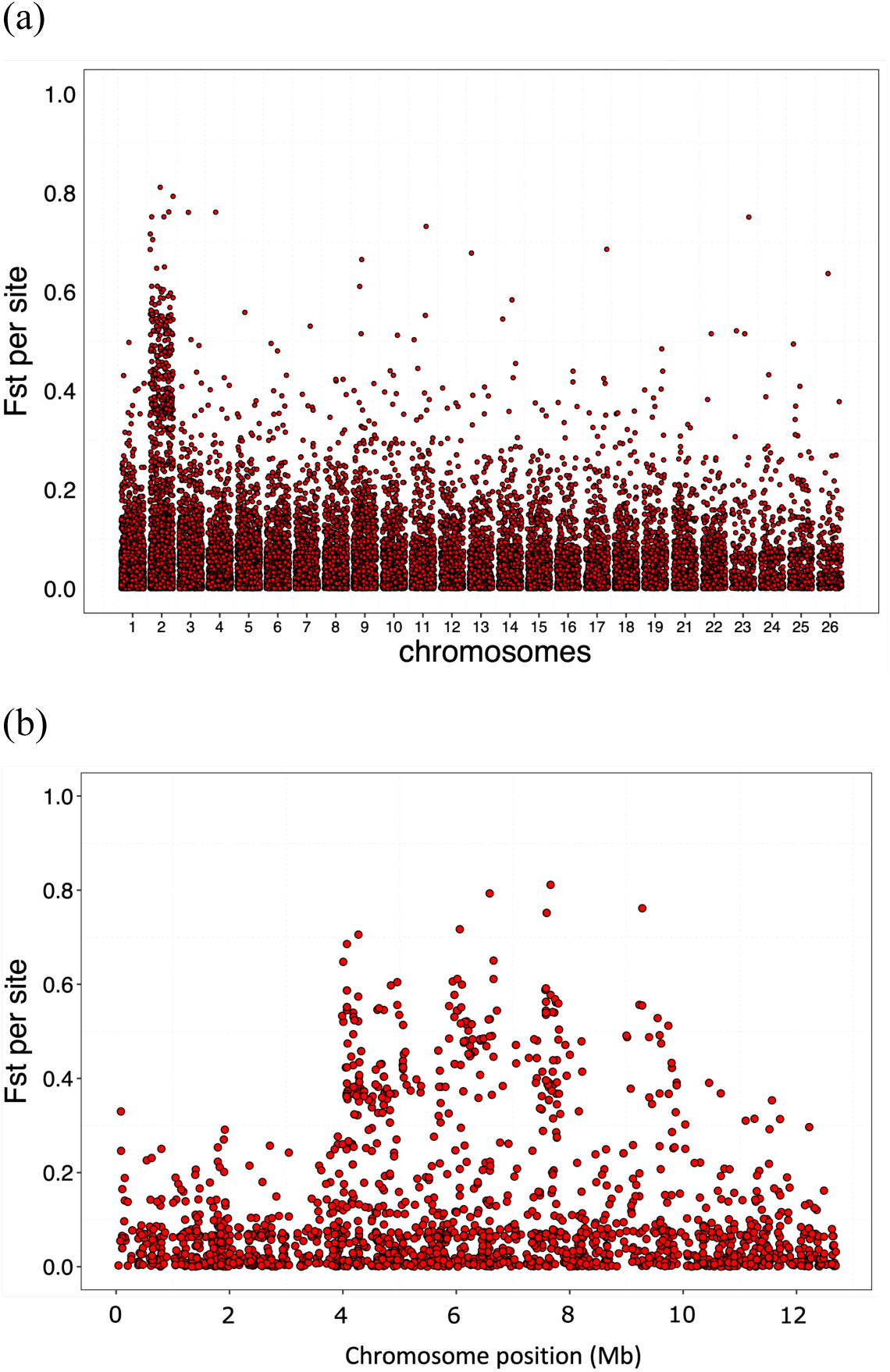
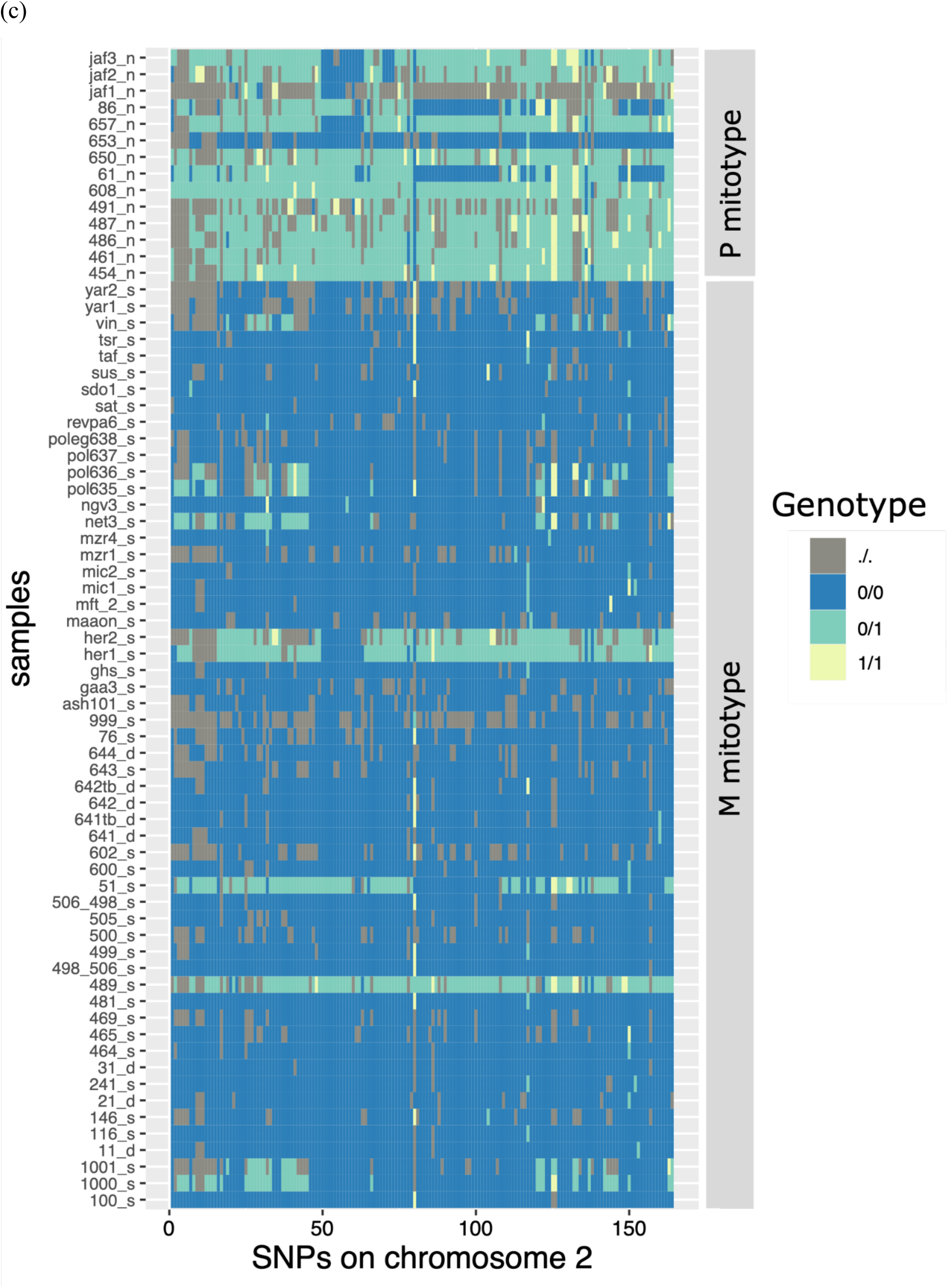
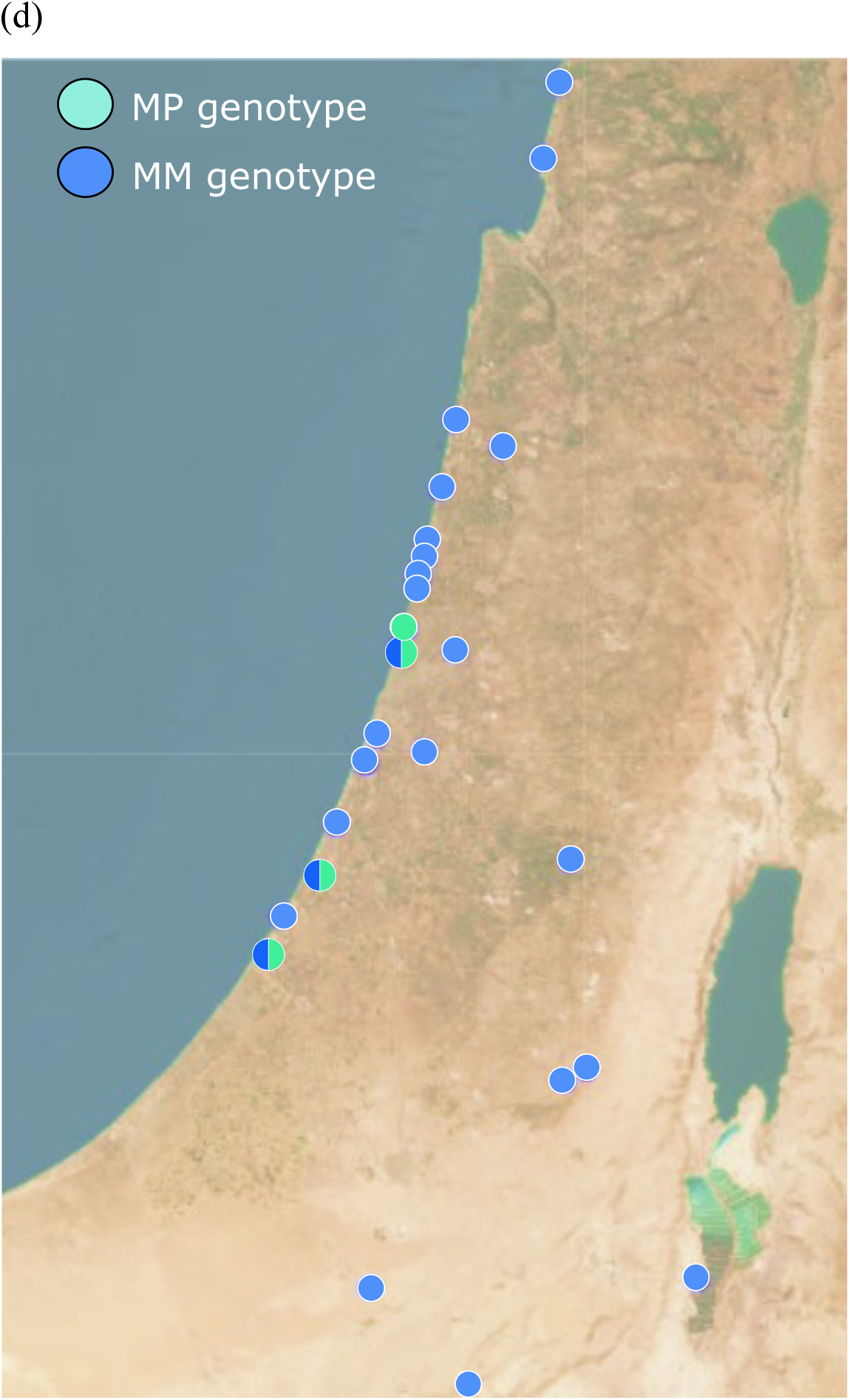
Genomic data of broad scale sampling of populations of *C. niger* across Israel from Reiner Brodetzki et al. (2019), showing the presence of the supergene on chromosome 2. Monogyne and polygyne samples were assigned based on the mitotype. (a) Genome-wide *FST* analysis using broad sampling of *C. niger* also shows a large number of high *FST* SNPs on chromosome 2. (b) The high *FST* region on chromosome 2 is consistent with our data from a single population. (c) Genotypes at the high *FST* SNPs show the association between mitotypes and supergene genotypes. Samples bearing the polygyne mitotype are mostly heterozygous MP and monogyne samples are homozygous MM at the supergene. (d) Distribution of supergene genotypes across Israel.

**Supplementary Figure S8:**
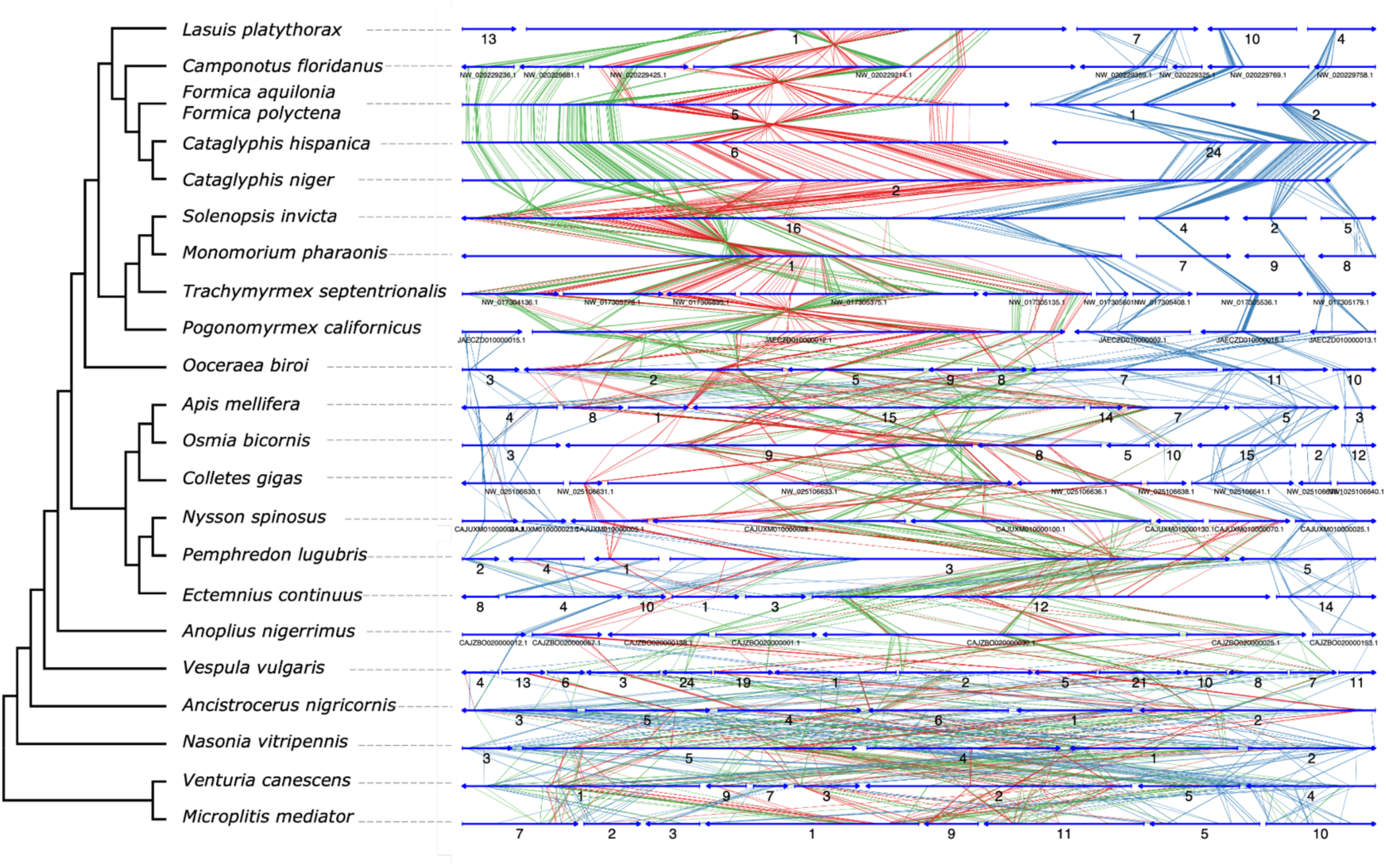
Synteny plot with 22 hymenopteran species based on one-to-one orthologous genes on the social chromosome of *Cataglyphis niger*.

**Supplementary Figure S9:**
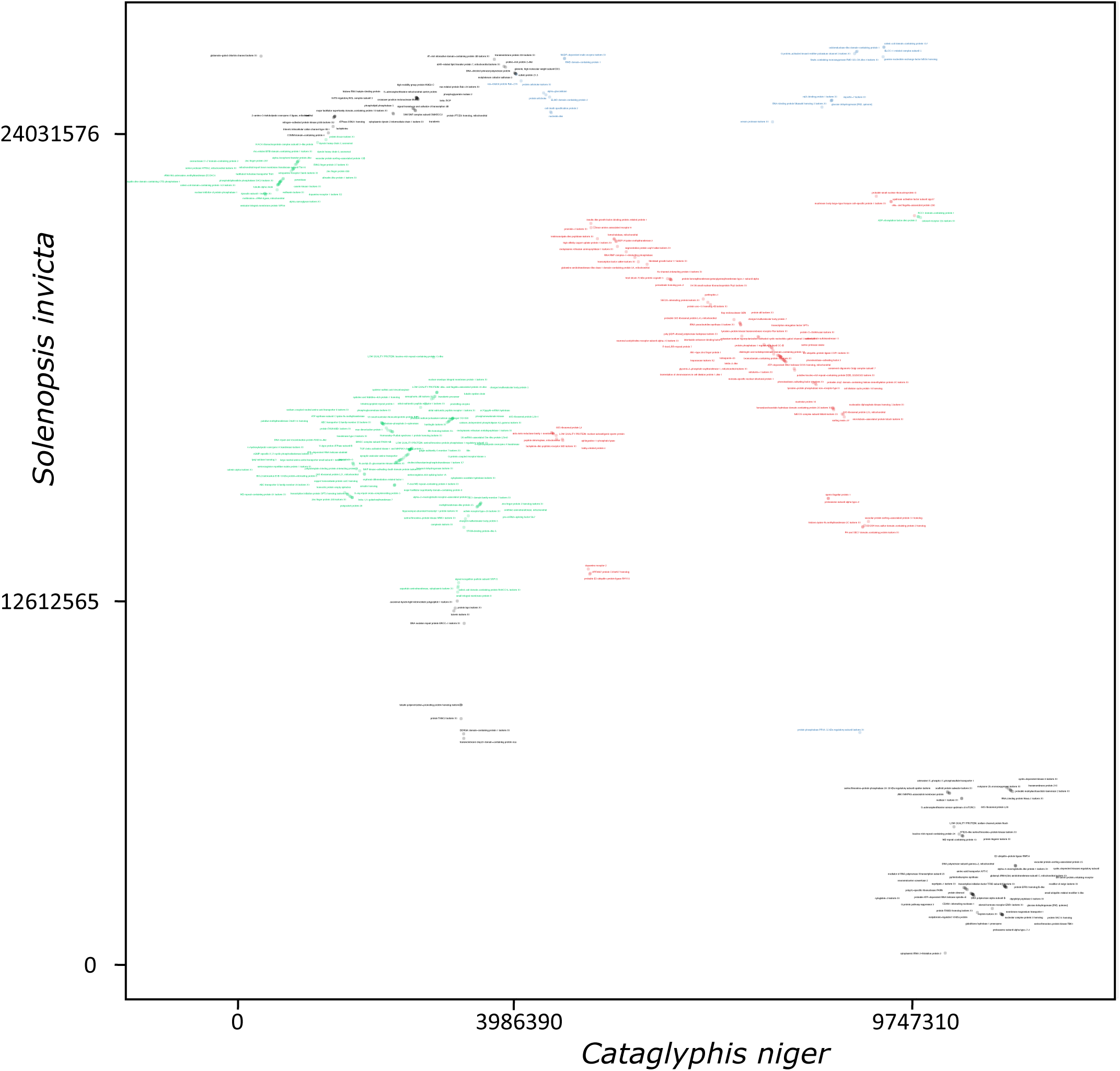
Dotplot of *Solenopsis* and *Cataglyphis* social chromosomes with gene names for one-to-one orthologs. Red dots indicate genes in the supergene region in both species.

**Supplementary Figure S10:**
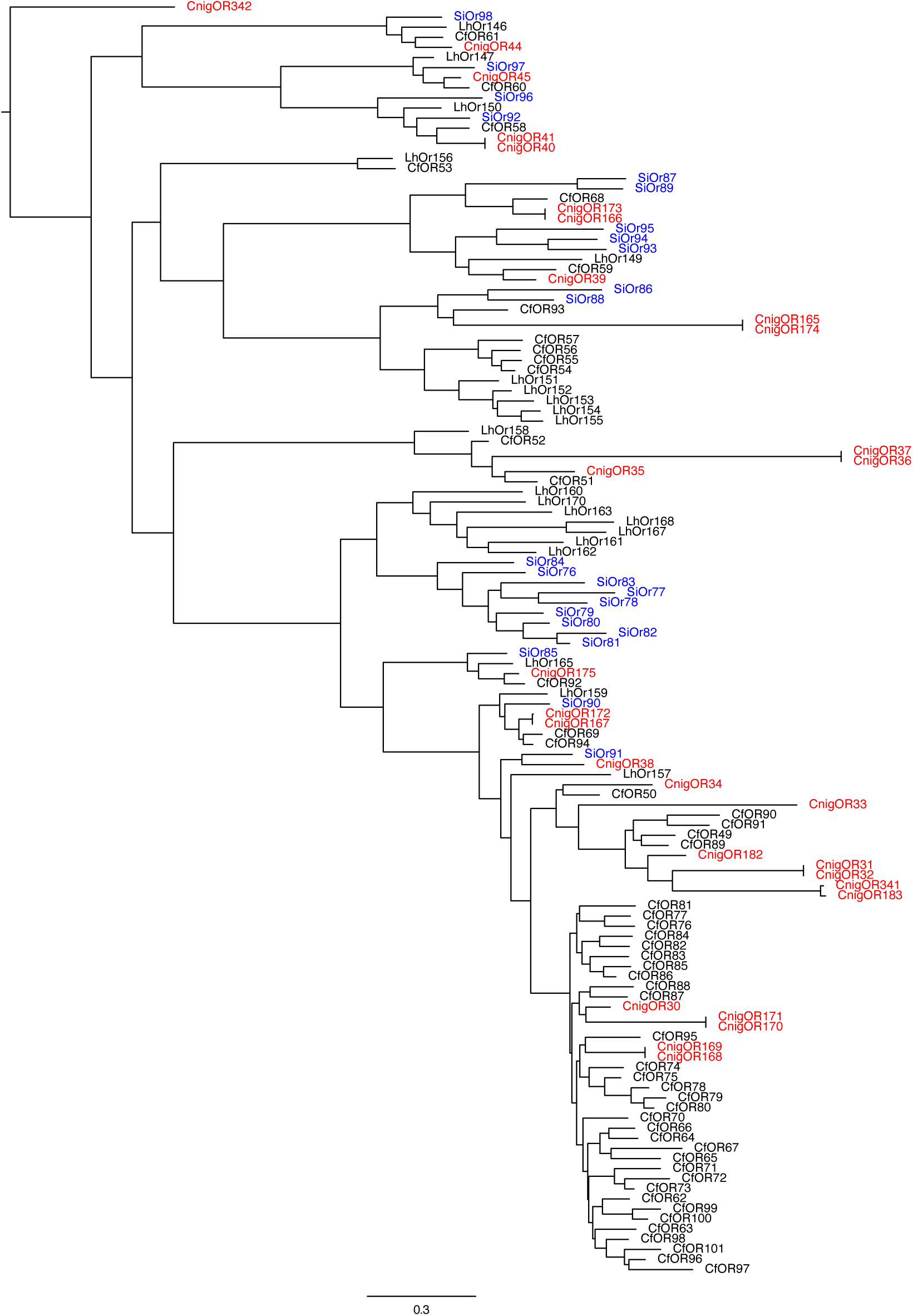
Subtree of the Odorant Receptor (OR) gene tree including genes in the *Cataglyphis* and *Solenopsis* social chromosomes. CnigOR = *Cataglyphis niger* OR; CfOR = *Camponotus floridanus* OR; SiOR = *Solenopsis invicta* OR; LhOR = *Linepithema humile* OR.

